# Rac1 deficiency reduces mitochondrial respiratory capacity, impairs fatty acid metabolism and causes muscle wasting

**DOI:** 10.64898/2026.06.14.732107

**Authors:** Lisbeth L. V. Møller, Steffen H. Raun, Emma Frank, Andreas B. Jordy, Nicoline R. Andersen, Anders Gudiksen, Zakarias Ogueboule, T. C. Phung Pham, Jessica L. Braun, Esben N. Poulsen, Jeffrey Molendijk, Anders Karlsen, Jakob Agergaard, Sean A. Newsom, Bryan C. Bergman, Niels Ørtenblad, Michael Kjær, Henriette Pilegaard, Bente Kiens, David E. James, Benjamin L. Parker, Joachim Nielsen, Steen Larsen, Erik A. Richter, Lykke Sylow

## Abstract

**Background:** The age-related progressive decline in skeletal muscle function is characterised by declining mitochondrial quality control and perturbed fatty acid metabolism, contributing to frailty and increased mortality. The actin cytoskeleton, a key structural component of skeletal muscle, has recently been implicated in mitochondrial anchoring and dynamics. However, the role of actin-regulating proteins, including the Rho GTPase Rac1, in mitochondrial function and age-associated metabolic and functional muscle deterioration remains undefined.

**Methods:** Skeletal muscle from mice with inducible muscle-specific deletion of Rac1 (Rac1 imKO) underwent unbiased mass spectrometry-based proteomic profiling. Mitochondrial morphology was assessed by transmission electron microscopy, and physiological parameters, including muscle mass and contraction-stimulated palmitate oxidation in isolated soleus muscle, were evaluated. Mitochondrial respiratory function was determined by high-resolution respirometry in permeabilised gastrocnemius skeletal muscle fibre bundles. Biochemically, muscular triacylglycerol (TG) content, mRNA (qPCR) and protein (immunoblotting) content were determined. In vastus lateralis muscle biopsies from healthy, untrained young (20-30 years) and old, sarcopenic (83-94 years) men, Rac1 and mitochondrial respiratory protein abundances were measured. A complementary human genetic association analysis was performed using the FinnGen dataset.

**Results:** Rac1 deficiency triggered muscle wasting in middle-aged mice (Gastrocnemius: -10%; Quadriceps: -7%). Preceding muscle wasting, gene set enrichment analysis identified enrichment in fatty acid metabolism and oxidative phosphorylation pathways, consistent with increased mitochondrial volume density in Rac1 imKO muscle (subsarcolemmal: +467%; intermyofibrillar: +166%). Despite mitochondrial expansion at this stage, Rac1 deficiency attenuated the increase in palmitate oxidation in response to muscle contraction (-62%). At the muscle-wasting stage, Rac1 imKO muscle exhibited reduced mitochondrial respiratory capacity (-25-32%). Additionally, the mitochondrial dysfunction was associated with an accumulation of muscle TG (+78%, p = 0.096) and upregulation of fatty acid transporter, CD36 protein content (+25%), indicative of altered fatty acid handling. In humans, Rac1 muscle protein content was increased in old, sarcopenic subjects compared to young (+41%), and negatively correlated with quadriceps cross-sectional area (CSA) (r = -0.475) and type II fibre CSA (r = -0.466). In old, sarcopenic muscle, Rac1 protein content correlated negatively with protein content of multiple mitochondrial respiratory complexes (CI: r = -0.690, CIV: r = -0.938, CV: r = -0.704). GWAS further identified associations between Rac1 SNP variants and lipid metabolic and muscle-wasting diseases.

**Conclusions:** Muscle Rac1 deficiency reduces mitochondrial respiratory capacity and metabolic flexibility through impaired fatty acid metabolism, leading to muscle wasting and highlighting a potential therapeutic target in age-related functional decline.

## Introduction

Skeletal muscle declines with age, manifesting as sarcopenia ^1^, a loss of muscle mass, strength and function that markedly increases morbidity and mortality ^2^. The progressive decrease in muscle function with age is characterised by declining mitochondrial quality control ^3^ and disrupted metabolism ^4,5^. Further, in older adults, muscle function is strongly associated with maximal mitochondrial phosphorylation capacity ^6^, yet the molecular mechanisms behind remain poorly understood. Defining these pathways is critical, as no clinically approved therapies exist to prevent or reverse sarcopenia, highlighting an opportunity for intervention.

An emerging player in myocellular function and mitochondrial biology is the Rho GTPase, Rac1, which controls the actin cytoskeleton dynamics in skeletal muscle cells ^7,8^. A crucial role for Rac1 in skeletal muscle insulin sensitivity by regulating the transport step of glucose into the muscle is well established ^7,9–11^, while the role of Rac1 in mitochondrial function, including fuel oxidation and energy production, and in age-related metabolic and functional muscle decline, remains unknown. Rac1 is predominantly localised to the cytosol and plasma membrane. Intriguingly, Rac1 can interact with the anti-apoptotic outer mitochondrial membrane protein Bcl-2 in non-muscle chronic myeloid leukaemia and Jurkat cells ^12^. In skeletal muscle, Rac1 might be implicated in mitochondrial regulation, as mice lacking the Rac1 downstream actin cytoskeleton-regulating targets, PAK1 and PAK2, develop megaconial mitochondria ^13^. The actin cytoskeleton is a key structural component of muscle, and an emerging new avenue of research is the interaction between mitochondrial anchoring and actin cytoskeleton dynamics. It has recently been proposed that actin cytoskeleton reorganisation is required for mitochondrial dynamics in non-muscle cells via a process where actin assembles on healthy mitochondria, aiding fission and blocking fusion^14^. After fission, actin disassembles, allowing rapid re-fusion and integration of the mitochondria into the mitochondrial network ^14^. This discovery and other studies ^15–19^ implicate the actin cytoskeleton in mitochondrial dynamics, suggesting that Rac1 may play a key role in mitochondrial function and fatty acid metabolism, yet this has never been demonstrated in skeletal muscle.

In human skeletal muscle, mitochondrial energetics associate with muscle quality ^6^. In rodents, gain- and loss-of-function studies have established mechanistic evidence and biological plausibility supporting a causal role for mitochondrial homeostasis in controlling muscle mass. For example, the ablation of Opa1 (Optic Atrophy 1), a mitochondrial dynamin-like GTPase that regulates mitochondrial inner membrane fusion, leads to ER stress, which signals via the unfolded protein response (UPR) and FoxOs, inducing a catabolic program of muscle loss ^20^. Also, both inhibition of mitochondrial fusion via muscle-specific ablation of MFN1 and MFN2 ^21,22^, and inhibition of mitochondrial fission in inducible muscle-specific knockout of Drp1 in mouse causes profound muscle atrophy ^23^. Thus, evidence ranging from human to preclinical mechanistic data points to mitochondria as key in age-related muscle decline and the maintenance of muscle mass and function. Hence, understanding mitochondrial regulation in preserving muscle mass and strength during ageing is crucial. While Rac1 has been implicated in exercise training-induced hypertrophy ^24^ and exercise-stimulated glucose uptake ^25–27^ in mouse muscle, the potential for Rac1 in age-related muscle wasting and mitochondrial biology in muscle has not been directly investigated. Interestingly, the development of sarcopenia is associated with alterations in actin cytoskeleton dynamics and fatty acid metabolism in human skeletal muscle ^4^. Thus, in this study, we hypothesised that Rac1 is implicated in muscle decline and wasting via alterations in actin dynamics, fatty acid metabolism, and mitochondrial dynamics. Our results reveal that Rac1 deficiency reduces mitochondrial respiratory capacity and metabolic flexibility through impaired fatty acid metabolism, leading to muscle wasting.

## Methods

### Ethical approval

Clinical experiments were approved by the Danish Regional Ethical Committees of the Capital Region approved the protocol (H-4-2013-068, H-2-2010-100), by the Danish Data Protection Agency (2007-58-0015, 2011-41-5965) and performed in accordance with the Helsinki Declaration II, and informed consent was obtained from all human research participants. Studies including human study participants are described in ^28–30^ and registered at https://ClinicalTrials.gov (NCT01997320, NCT01252381). All mouse experiments complied with the European Convention for the protection of vertebrate animals used for experimental and other scientific purposes (No. 123, Strasbourg, France, 1985; EU Directive 2010/63/EU for animal experiments) and were approved by the Danish Animal Experiments Inspectorate (License: 2015-15-0201-00477).

### Human skeletal muscle biopsies

Vastus lateralis muscle biopsies from old, sarcopenic men (age 83-94 years) were obtained as described ^29,30^. Only baseline biopsies from male participants from the original studies were investigated. In addition, baseline biopsies from healthy, sedentary young males (aged 20–30 years) from another study ^28^ were included as the young control group for comparison between young men and old, sarcopenic men. Based on the available biopsy material, biopsies from 10 young participants and 9 old, sarcopenic participants were included in the analyses. Quadriceps and fibre type cross-sectional area (CSA) have previously been published ^28–30^.

### Mouse Studies

All mice were maintained on a 12:12-h light–dark cycle and housed at 22 °C (with allowed fluctuation of ±2 °C) with nesting material. The mice were group-housed. The mice received a standard rodent chow diet (Altromin no. 1320; Brogaarden, Denmark) and water ad libitum. Male and female tetracycline-inducible muscle-specific Rac1 knockout (imKO) mice were used for experiments, except for the subcellular fractionations, where muscles from female C57Bl/6N wild-type spare mice from breeding of SLIRP knockout mice ^31^ were analysed.

### Tetracycline-inducible muscle-specific Rac1 imKO mice

The generation of the tetracycline-inducible muscle-specific Rac1 imKO mice has been described elsewhere ^25^. Mice were backcrossed until N5 (96.9% congenic) on a C57BL/6JBomTac background. Transgenic mice were littermates from the breeding of heterozygous Cre and Rac1 *fl/fl* transgenic mice. Both males and females were used for experiments conducted in adult mice, whereas only male, middle-aged mice were used as indicated in the text and figures. Knockout was obtained by adding the tetracycline analogue doxycycline (1 g/L; Sigma-Aldrich) to the drinking water for 21 days, after which it was switched to normal tap water. Mice were used for experiments after a wash-out period of at least 3 weeks. For experiments conducted in adult mice (4.4 ± 0.8 months old, mean ± SD), doxycycline treatment was initiated at age 2.5 ± 0.6 months. Unless otherwise stated, fed adult mice were anaesthetised (6 mg pentobarbital sodium 100 g^−1^ body weight i.p.) and skeletal muscles were excised and quickly frozen in liquid nitrogen and stored at -80°C until processing. The mice were euthanised by cervical dislocation after muscle isolation. For experiments conducted in middle-aged, male mice (12.7 ± 0.3 months old), doxycycline treatment was initiated at age 2.9 ± 0.7 months. Middle-aged mice were re-treated with doxycycline for three days every third month. One-hour fasted middle-aged mice were anaesthetised (6 mg pentobarbital sodium 100 g^−1^ body weight i.p.) followed by cervical dislocation. Skeletal muscles were excised and quickly frozen in liquid nitrogen and stored at -80°C until processing.

### Mouse genotyping

Mouse genotyping was performed as previously described ^32^. In brief, an ear punch was digested overnight in DirectPCR Lysis Reagent (#101-T, Viagen Biotech) with 0.2 mg mL^-1^ Proteinase K (# 03115828001, Roche) at 55°C, followed by 1 hour at 85°C. After spin at 1000 g for 5 min, the supernatant was diluted 10X in TE pH 8.0 with 50 pg mL^-1^ Quinoline Yellow. Real-time quantitative PCR was performed using a MX3005P real-time PCR machine using Quantitect SYBR Green mastermix (Qiagen), primers listed in Table S1 and 5 pg mL^-1^ Xylene Cyanol. The reactions were furthermore spiked with a heterozygote sample as a positive control. The Ct values were used to assess allele presence by comparison to the no DNA controls (spike values). Amplification efficiency in the individual reactions was estimated by the sigmoid method of Liu and Saint ^33^ to ensure that the Ct’s could be compared within primer sets. The genotype was later verified by immunoblotting on muscle tissue.

### Morphological analyses

Body composition was analysed using magnetic resonance imaging (EchoMRI-4in1TM, Echo Medical System LLC, Texas, USA).

### Mass spectrometry proteomic analysis

#### Sample preparation

##### Adult mice

Mice were euthanised by cervical dislocation, and gastrocnemius muscles were excised and quickly frozen in liquid nitrogen and stored at -80°C until processing. Samples were lysed in 6 M urea, 2 M thiourea, 25 mM triethylammonium bicarbonate (TEAB), pH 7.9, containing phosphatase and protease inhibitor cocktail by tip-probe sonication (2 × 15 s) on ice. The lysates were centrifuged at 17,000 x g, 15 min, 4°C and the supernatant precipitated with six volumes of acetone overnight at −20°C. Protein pellets were resuspended in 6 M urea, 2 M thiourea, 25 mM TEAB, pH 7.9 and quantified by Qubit fluorescence (Invitrogen by LifeTechnologies, Carlsbad, CA, USA). Concentrations were normalised to 1 mg mL^-1^ and protein reduced with 10 mM dithiothreitol for 60 min at 25°C, followed by alkylation with 25 mM iodoacetamide for 30 min at 25°C in the dark. The reaction was quenched to a final concentration of 20 mM dithiothreitol and digested with Lys-C (WakoPure Chemical Industries, Osaka, Japan) at 1:50 enzyme-to-substrate ratio for 2 h at 25°C. The mixture was diluted 5-fold with 25 mM TEAB and digested with sequencing-grade trypsin at 1:50 enzyme-to-substrate ratio for 12 h at 30°C. The peptide mixture was acidified to a final concentration of 2% formic acid, 0.1% trifuoroacetic acid (TFA) and centrifuged at 16,000 g for 15 min. Peptides were desalted using hydrophilic lipophilic balance–solid phase extraction (HLB-SPE) cartridges (Waters Corp., Milford, MA, USA) followed by elution with 50% acetonitrile, 0.1%TFA, and dried by vacuum centrifugation. Peptides were resuspended in 30 μL of 100 mM TEAB, quantified by Qubit fluorescence and normalised to 10 mg mL^-1^. Peptides were labelled with 10-plex Tandem Mass Tags (TMT) in a final concentration of 50% acetonitrile for 1.5 h at room temperature and deacylated with 0.3% hydroxylamine. The labelled peptides were acidified to 0.1% TFA, pooled and dried to approximately 50 μL by vacuum centrifugation. Peptides were resuspended in 0.1% TFA and desalted with HLB-SPE cartridges, dried and stored at -30°C until analysis. Peptides were fractionated into 12 fractions on an in-house packed TSKgel-amide (Tosoh Bioscience, Griesheim, Germany) hydrophilic interaction chromatography (HILIC) column as described previously ^34^.

##### Middle-aged mice

Tissue samples were lysed by the addition of EasyPEP lysis buffer (Thermo Fisher Scientific) supplemented with nuclease using the BeatBox tissue disruptor (Preomics) for 10 minutes at standard setting. Samples were incubated for 10 min at 95°C, followed by additional 10 min of BeatBox. Protein concentrations were measured (Pierce BCA Protein Assay Kit, Thermo), and 100 µg aliquots were prepared. Samples were reduced with 5 mM (final concentration) of TCEP for 15 min at 55°C, alkylated with 20 mM (final concentration) CAA for 30 minutes at RT, and digested by adding Trypsin/LysC at 1:50 enzyme/protein ratio. Peptides were acidified with 10% formic acid (FA), desalted using a Thermo desalting plate and dried down using vacuum centrifugation. Peptides were resuspended in A* (0.1% TFA, 2% ACN) and stored at -20 °C prior to LC-MS analysis.

#### Mass spectrometry

##### Adult mice

Peptides were resuspended in 2% acetonitrile, 0.5% acetic acid and loaded onto a 50 cm × 75 μm inner diameter column packed in-house with 1.9 μm C18AQ particles (Dr. Maisch HPLC GmbH, Ammerbuch-Entringen, Germany) using a Dionex UHPLC. Peptides were separated using a linear gradient of 5–30% Buffer B over 120 min at 250 nl min^−1^ (Buffer A = 0.5% acetic acid; Buffer B = 80% acetonitrile, 0.5% acetic acid). The column was maintained at 50°Cusing a PRSO-V1 ion-source (Sonation) coupled directly to a Q-Exactive Plus mass spectrometer (MS). A first full-scan MS was measured at 70,000 resolution at 200 m/z (300–1550 m z^−1^ ; 100 ms injection time; 3e6 automatic gain control (AGC) target) followed by isolation of up to 20 of the most abundant precursor ions for MS/MS (1.2 m/z isolation; 8.3e5 intensity threshold; 30.0 normalized collision energy; 35,000 resolution at 200 m/z; 120 ms injection time; 2e5 AGC target).

##### Middle-aged mice

Peptides were separated on an Aurora (Gen3) 25 cm, 75 μM ID column packed with C18 beads (1.6 μm) (IonOpticks) using a Vanquish Neo (Thermo Fisher Scientific) UHPLC. Peptide separation was performed using a 90 min stepped gradient of 2-17% solvent B (0.1% formic acid in acetonitrile) over 56 min, 17-25% solvent B over 21 min, 25-35% solvent B over 13 min, using a constant flow rate of 400 nL/min. Column temperature was maintained at 50 °C. Upon elution, peptides were injected via a CaptiveSpray2 source and 20-μm emitter into a timsTOF Pro2 mass spectrometer (Bruker) operated in diaPASEF mode. MS data were collected over a 100-1700 m/z range. The standard ’long-gradient’ diaPASEF method was used, which included 16 diaPASEF scans with two 26 Da windows overlapping by 1 Da per ramp, a total mass range of 400-1201 Da, and a mobility range of 1.6-0.6 1/K0. The collision energy was decreased linearly from 59 eV at 1/K0 = 1.6 to 20 eV at 1/K0 = 0.6 Vs cm-2. Both accumulation time and PASEF ramp time were set to 100 ms. The total cycle time was 1.8 s.

#### Data processing and bioinformatics

##### Adult mice

Data were processed using MaxQuant v1.6.12.0^35^ and searched with Andromeda ^36^ against the mouse UniProt database (January 2022; 55,004 entries). The data were searched with variable modification of methionine oxidation and cysteine carbamidomethylation as a fixed modification. The precursor-ion mass tolerance was set to 20 p.p.m. and 7 p.p.m. for first and second searches, respectively, and product-ion mass tolerance set to 0.02 Da. All results were filtered to 1% false discovery rates (FDRs). All data were normalised to the median of each replicate. Significantly regulated proteins were determined using t-tests corrected for multiple testing using Benjamini–Hochberg in the Perseus Software Package ^37^.

##### Middle-aged mice

MS files were processed using Spectronaut version 20.1 (Biognosys) in direct DIA search mode with the default workflow. Briefly, UniprotKB UP0000000589 (Taxa ID: 10090) Mus musculus FASTA database was used. Carbaminomethylation of cysteine was set as a fixed modification, whereas the oxidation of methionine and the acetylation of the protein N-terminus were set as variable modifications. The maximum number of missed cleavages was 2, and the minimum peptide amino acid length was 7. The false discovery rate for PSM, peptide, and protein groups was set to 0.01. All statistical analyses were performed using in-house developed python code, based on the automated analysis pipeline of the Clinical Knowledge Graph ^38^. Intensity values were log2-transformed, and features with fewer than 70% of valid values in at least one group were removed. Remaining missing values were replaced by mixed imputation, where kNN and MinProb (width=0.3 and shift=1.8) methods are used for values missing at random (MAR) and values missing not at random (MNAR), respectively ^39^. MAR is defined when a minimum 60% of the samples within a given group have an existing value. Differentially expressed features were identified by statistical paired and unpaired t-tests, and Benjamini-Hochberg correction for multiple hypothesis testing with False Discovery Rate (FDR) threshold 0.05, and fold-change of 2.

### Transmission electron microscopy

A longitudinal section of the quadriceps muscle tissue was fixed by immersion in 2.5% glutaraldehyde in 0.1 M sodium cacodylate buffer (pH 7.3) and stored at 4°C. The fixed specimen was rinsed four times in 0.1 M sodium cacodylate buffer, before being post-fixed with 1% osmium tetroxide (OsO4) in 0.1 M sodium cacodylate buffer for 90 min at 4°C. The muscles were then rinsed twice in 0.1 M sodium cacodylate buffer at 4°C, dehydrated through a graded series of alcohol at 4–20°C, infiltrated with graded mixtures of propylene oxide and Epon at 20°C, and embedded in 100% Epon at 30°C, as previously described ^40^. Ultra-thin (60 nm) longitudinal sections were cut (using an Ultracut UCT ultramicrotome; Leica Microsystems, Wetzlar, Germany) in two depths (separated by 150 μm) and contrasted with uranyl acetate and lead citrate. Sections were examined and images captured with a pre-calibrated CM100 TEM (Philips, Eindhoven, The Netherlands) operated at 100 kV, and equipped with a Veleta camera (Olympus, Hamburg, Germany) and the iTem software package at a resolution of 4008 × 2664 pixels.

Except for one Rac1 imKO specimen, six longitudinally oriented fibres were photographed and from each fiber, 24 images were obtained at x13,500 magnification. The imaging was performed in a randomized systematic order including 12 images from the subsarcolemmal (SS) region, and 12 from both the superficial and central region of the intermyofibrillar (IMF) space. The volumetric content of intermyofibrillar and subsarcolemmal mitochondria was estimated by point counting ^41^ using a grid size of 460 nm generating 256 points per image and expressed per myofibrillar space and fibre surface area, respectively. The mitochondrial cristae surface area per mitochondrial volume was estimated by counting intersections on test lines ^42^ using a grid size of 90 nm. The Z-disc width was measured once on all IMF images. The quantification of other mitochondrial morphology measures was done using the Radius EM Imaging Software (Emsis GmbH, Radius 2.0). The images were analysed by a set-up blinded investigator.

### Lipid profile

Fractions collected containing cytosolic and membrane compartments were brought to 3 mL total volume, then 1.5 mL MeOH and 5 mL MTBE were added. Then, an internal standard cocktail was added, and samples were lipid-extracted. The top fraction containing the lipids was saved, and the lipid extraction was repeated. Triacylglycerol (TG), diacylglycerol (DAG), acylcarnitine and sphingolipid species were analysed by an Agilent 1100 HPLC connected to an API 2000 triple quadrupole mass spectrometer as previously described ^43^. The 1,3- and 1,2-DAG isomers were separated chromatographically using a Hilic 2.1 micron, 3x100 mm column. Concentration was determined by comparing ratios of unknowns to class-specific internal standards and compared to standard curves representing the majority of lipid species run with each sample set.

### Metabolomics

#### Sample preparation

Mice were acclimatised to treadmill running, and maximal running capacity speed was determined as previously described ^27,44^. Fed mice ran on a treadmill at 70-75% of the average maximal running speed (10° incline). After 30 minutes, mice were euthanised by cervical dislocation and gastrocnemius muscles were excised and quickly frozen in liquid nitrogen and stored at -80°C until processing. Untargeted metabolomic profiling was performed at Metabolon, Inc. (Durham, NC). Samples were prepared using the automated MicroLab STAR® system (Hamilton Company). Proteins were precipitated with methanol under vigorous shaking for 2 min (Glen Mills GenoGrinder 2000) followed by centrifugation. The resulting extract was divided into four fractions: one for analysis by UPLC-MS/MS with positive ion mode electrospray ionization, one for analysis by UPLC-MS/MS with negative ion mode electrospray ionization, one for LC polar platform, and one for analysis by GC-MS. Samples were placed briefly on a TurboVap® (Zymark) to remove the organic solvent.

#### Ultrahigh Performance Liquid Chromatography-Tandem Mass Spectroscopy (UPLC-MS/MS)

The LC/MS portion of the platform was based on a Waters ACQUITY ultra-performance liquid chromatography (UPLC) and a Thermo Scientific Q-Exactive high resolution/accurate mass spectrometer interfaced with a heated electrospray ionization (HESI-II) source and Orbitrap mass analyser operated at 35,000 mass resolution. The samples were stored overnight under nitrogen before preparation for analysis. The sample extract was dried, then reconstituted in acidic or basic LC-compatible solvents, each of which contained 8 or more injection standards at fixed concentrations to ensure injection and chromatographic consistency. One aliquot was analysed using acidic positive ion optimized conditions and the other using basic negative ion optimized conditions in two independent injections using separate dedicated columns (Waters UPLC BEH C18-2.1x100 mm, 1.7 µm). Extracts reconstituted in acidic conditions were gradient eluted from a C18 column using water and methanol containing 0.1% formic acid. The basic extracts were similarly eluted from C18 using methanol and water, however with 6.5mM Ammonium Bicarbonate. The third aliquot was analysed via negative ionization following elution from a HILIC column (Waters UPLC BEH Amide 2.1x150 mm, 1.7 µm) using a gradient consisting of water and acetonitrile with 10mM Ammonium Formate. The MS analysis alternated between MS and data-dependent MS2 scans using dynamic exclusion, and the scan range was from 80-1000 m/z.

#### Gas Chromatography-Mass Spectroscopy (GC-MS)

The samples destined for analysis by GC-MS were dried under vacuum for a minimum of 18 h prior to being derivatized under dried nitrogen using bistrimethyl-silyltrifluoroacetamide. Derivatized samples were separated on a 5% diphenyl / 95% dimethyl polysiloxane fused silica column (20 m x 0.18 mm ID; 0.18 um film thickness) with helium as carrier gas and a temperature ramp from 60° to 340°C in a 17.5 min period. Samples were analysed on a Thermo-Finnigan Trace DSQ fast-scanning single-quadrupole mass spectrometer using electron impact ionization (EI) and operated at unit mass resolving power. The scan range was from 50–750 m/z.

#### Bioinformatics

Raw data was extracted and peak-identified using Metabolon’s hardware and software. Compounds were identified by comparison to library entries of purified standards or recurrent unknown entities. Biochemical identifications are based on three criteria: retention index within a narrow retention time/index (RI) window of the proposed identification, accurate mass match to the library +/- 0.005 amu, and the MS/MS forward and reverse scores between the experimental data and authentic standards. Peaks were quantified using area-under-the-curve. Values were normalised in terms of raw area counts, and each biochemical was rescaled to set the median equal to 1. Missing values were imputed with the minimum observed value for each compound. Following log transformation, a Welch’s two-sample t-test was used to identify biochemicals that differed significantly between experimental groups.

### Fatty acid oxidation in isolated soleus muscles

#### Electrical stimulation in vitro

Exogenous palmitate oxidation in isolated soleus muscles of Rac1 imKO mice was analysed using procedures described previously ^45^. Soleus muscle were dissected from anaesthetised (6 mg pentobarbital sodium 100 g^−1^ body weight i.p.) male mice and suspended at resting tension (4–5 mN) mounted to a force transducer in 15 ml incubation reservoirs (Radnoti, CA, USA) in Krebs-Henseleit Ringer buffer pH 7.4 containing 5 mM glucose, 2% fatty acid free BSA and 0.5 mM palmitic acid at 32°C oxygenated prior to use. Palmitic acid was dissolved in 96% ethanol, and a small volume was added to the buffer (<1% total buffer volume) to achieve the desired palmitate concentration. The mice were euthanised by cervical dislocation after muscle isolation. After a 20-minute pre-incubation, the incubation buffer was replaced with the same buffer described above supplemented with 0.5 μCi mL^-1^ of [1-^14^C] palmitate (Perkin Elmer, MA, USA). Fatty acid oxidation was measured in resting or contracting soleus muscle over 25 min. Contractions were induced by electrical stimulation every 3.3 sec with 600 msec trains of 2 msec pulses delivered at 30 Hz (60V). After the contraction protocol, muscles were removed and snap-frozen in liquid nitrogen and stored at -80°C. The incubation medium was collected, and gaseous ^14^CO_2_ was liberated with 1 M acetic acid and trapped with vials containing 0.4 ml benzethonium hydroxide. Radioactivity in trapped ^14^CO_2_ was determined by liquid scintillation counting using Ultima Gold (Perkin Elmer, MA, USA) – representing the complete fatty acid oxidation. Frozen muscle strips were quickly trimmed of visible connective tissue and surrounding adipose tissue and weighed. Muscles were cut in half, and lipids were extracted from one half by homogenization in chloroform:methanol (2:1) and centrifuged for 10 min at 2,714 x g. Then, H_2_O was added, samples gently vortexed and centrifuged for 10 min at 2,714 x g again. The aqueous phase was subjected to liquid scintillation counting, representing counts trapped in isotonic exchange (acid-soluble metabolites). Palmitate oxidation was determined as CO_2_ production (complete fatty acid oxidation) and acid-soluble metabolites (representing incomplete fatty acid oxidation); thus, counts obtained within the aqueous phase were combined with the counts from trapped ^14^CO_2_ to calculate rates of fatty acid oxidation.

#### Fatty acid incorporation into TAG and DAG

Fatty acid incorporation into TAG and DAG was measured using procedures described previously ^46^. The chloroform phase, which contains the total lipids extracted from muscle, was evaporated under a stream of N_2_ and redissolved in chloroform. Each sample was spotted on TLC silica-gel coated plates and the lipids resolved in petroleum ether:diethyl ether:acetic acid (120:25:1.5) for 50 min placed in a sealed tank. Plates were air-dried and dipped in 10% copper sulfate pentahydrate and 8% phosphoric acid solution at 120°C for 15 minutes. The silica-gel coated plates were visualised by long-wave UV-light, TAG and DAG bands marked and scraped into scintillation vials for liquid scintillation. Palmitate uptake was determined as the sum of palmitate oxidation, palmitate incorporation into TAG and palmitate incorporation into DAG.

### Electrical stimulation *in situ*

After the soleus muscles were used for fatty acid metabolism experiments, the skin was removed from the right hindlimb, and needles were inserted at the sciatic nerve and at the tibio-patellar ligament. The quadriceps muscle was electrically stimulated to contract for 25 min. Contractions were induced by electrical stimulation every 3.3 sec with 600 msec trains of 2 msec pulses delivered at 30 Hz (5-6V). The contralateral leg served as a resting control.

### Measurement of muscle triacylglycerol (TG)

Biochemical measurements of muscle TG were performed as described previously ^47^. In short, freeze-dried and dissected muscle tissue or wet weight muscle tissue, as indicated, was incubated overnight in tetraethylammonium hydroxide. Next, 3 M perchloric acid was added, and the samples were centrifuged at 22°C for 10 min at 3,000 g before being neutralised with 2 M KHCO_3_. The content of glycerol was determined fluorometrically using ABX Penta c400 (Horiba ABX SAS).

### Measurement of muscle glycogen concentration

Muscle glycogen content was determined as glycosyl units after acid hydrolysis, as previously described ^47,48^. In short, freeze-dried and dissected muscle tissue was incubated for 2 h at 98°C with 1N HCl. Glycogen concentration as glycosyl units was determined fluorometrically using ABX Penta c400 (Horiba ABX SAS).

### Mitochondrial respiration

In a subset of the middle-aged mice, mitochondrial respiratory capacity was measured in permeabilised gastrocnemius skeletal muscle fibres as previously described ^49,50^. In brief, the gastrocnemius muscle was rinsed free of fat and connective tissue and separated into small fibre bundles. Fibre bundles were permeabilised with saponin (50 µg mL^-1^) in ice-cold BIOPS buffer for 30 min, followed by a 20 min wash in MiR05 buffer on ice. Mitochondrial respiration was measured in duplicate under hyperoxic conditions at 37 °C using high-resolution respirometry (Oxygraph-2k, Oroboros Instruments, Austria). The following protocol was applied: Leak respiration was assessed by the addition of malate (2 mM) and pyruvate (5 mM), followed by adding two different concentrations of ADP (0.25 mM and 5 mM) to assess state 3 respiration. 10 µM Cytochrome C was added to test the integrity of the outer mitochondrial membrane, and then 10 mM Glutamate was added to measure complex I-linked maximal state 3 respiratory capacity, followed by 10 mM succinate to measure complex I + II-linked maximal state 3 respiratory capacity. Finally, 2.5 µM Antimycin A was added to inhibit complex III in the electron transport chain for the assessment of non-mitochondrial background respiration.

### Citrate synthase (CS) activity

In the same subset of middle-aged mice chosen for mitochondrial respiration, CS activity was measured on Cobas 6000 (C 501, Roche Diagnostics) as described previously ^51^. The muscle tissue was homogenised for 2 minutes using a Tissuelyser II (Qiagen, Germany) in an assay-specific buffer (0.3 M K_2_HPO_4_, 0.05% (v/v) bovine serum albumin (BSA), pH 7.7) before adding Triton (0.1% (v/v)). The homogenate was diluted (0.33 mM acetyl-CoA, 0.6 mM oxaloacetate, 0.157 mM DTNB, 39 mM Tris-HCl (pH 8.0)) and then measured spectrophotometrically (415 nm) ^51,52^.

### Subcellular fractionation

The subcellular fractionation assay for frozen muscle was adapted ^53,54^ and performed on ice or at 4 °C, where applicable, as previously described ^55^. Frozen quadriceps muscle tissue was thawed in room temperature isolation buffer solution, herein referred to as ISO buffer, containing 880 mM sucrose, 20 mM HEPES (pH 7.4), 50 mM NaCl, 5 mM MgCl2, 5 mM EGTA, 20 mM β-glycerophosphate, 10 mM NaF, 2 mM phenylmethylsulfonyl fluoride, 1 mM EDTA (pH 8.0), 1 mM EGTA (pH 8.0), 2 mM Na_3_VO_4_, leupeptin (10 μg mL^-1^), aprotinin (10 μg mL^-1^), and 3 mM benzamidine. Room temperature ISO buffer was replaced with ice-cold ISO buffer, and muscle was minced finely with a scissor, followed by homogenization for 2 min at 17.5 Hz using a TissueLyser II bead mill (QIAGEN, USA). The homogenate was then rotated end-over-end for 30 min and centrifuged at 1000 g for 10 min. The resulting supernatant was used to prepare the cytosolic and mitochondrial fractions.

For the cytosolic and mitochondrial fractions, the supernatant fraction was centrifuged twice for 10 min at 1000 g. The resulting pellets were discarded, and the supernatant was centrifuged for 20 min at 20,000 g. From here, the resulting supernatant and pellet were used to obtain the cytosolic and mitochondrial fractions, respectively. For the cytosolic fraction, the supernatant was centrifuged twice for 20 min at 20,000 g and the resulting pellets were discarded, followed by first an ultracentrifugation a 100 000 g for 1 h, then a wash and ultracentrifugation at 100 000 g for 45 min in ISO buffer. The final supernatant contained the cytosolic fraction. For the mitochondrial fraction, the pellet was resuspended in ISO buffer, centrifuged twice for 20 min at 20 000 g and the resulting supernatants were discarded. The resulting pellet was resuspended in 75 μL mitochondrial lysis buffer [50 mM Tris HCl (pH 6.8), 1 mM EDTA, 0.5% Triton X-100, 20 mM β-glycerophosphate, 10 mM NaF, 2 mM phenylmethylsulfonyl fluoride, 1 mM EDTA (pH 8.0), 1 mM EGTA (pH 8.0), 2 mM Na3VO4, leupeptin (10 μg mL^-1^), aprotinin (10 μg mL^-1^), and 3 mM benzamidine], sonicated 3X 10 seconds and rotated end-over-end for 20 min. The final suspension contained the mitochondrial fraction.

### RNA extraction and quantitative real-time PCR

Total RNA was extracted from the gastrocnemius muscle using QIAzol lysis reagent (Qiagen) and RNeasy Mini Kit (Qiagen). For RT–qPCR expression analysis, 400 ng of RNA was reverse transcribed using the High-Capacity cDNA Reverse Transcription Kit with RNase Inhibitor (Applied Biosystems) as per the manufacturer’s protocol. The qPCR was performed in ViiA 7 Real-Time PCR System using QuantStudio Real-Time PCR software (Applied Biosystems, Waltham, MA), using TaqMan™ Universal Master Mix II, with UNG (Applied Biosystems). Cycle threshold (Ct) was converted to a relative amount using a standard curve derived from a serial dilution of a representative pooled sample run together with the samples of interest. 18S rRNA was used for normalisation of target mRNA levels. TaqMan gene expression assays used for mRNA levels shown in Table S2.

### Protein extraction

Muscle tissue was crushed in liquid nitrogen prior to homogenization. All tissues were homogenized 2 x 30 sec at 30 Hz using a Tissuelyser II (Qiagen, USA) in ice-cold homogenization buffer (10% (v/v) Glycerol, 1% (v/v) NP-40, 20 mM Na-pyrophosphate, 150 mM NaCl, 50 mM HEPES (pH 7.5), 20 mM β-glycerophosphate, 10 mM NaF, 2 mM PMSF, 1 mM EDTA (pH 8.0), 1 mM EGTA (pH 8.0), 2 mM Na3VO4, 10 μg mL^-1^ Leupeptin, 10 μg mL^-1^ Aprotinin, 3 mM Benzamidine). After rotation end-over-end for 30 min at 4°C, lysate supernatants were collected by centrifugation (10,854 x g) for 15 min at 4°C.

### Immunoblotting

Lysate protein concentration was determined using the bicinchoninic acid method using BSA standards and bicinchoninic acid assay reagents (Pierce). Immunoblotting samples were prepared in 6X sample buffer (340 mM Tris (pH 6.8), 225 mM DTT, 11% (w/v) SDS, 20% (v/v) Glycerol, 0.05% (w/v) Bromphenol blue). Protein phosphorylation (p) and total protein content were determined by standard immunoblotting technique, loading equal amounts of protein. Each antibody was optimised to ensure that the protein amount loaded was within a linear signal-to-protein range. The polyvinylidene difluoride membrane (Immobilon Transfer Membrane; Millipore) was blocked in Tris-Buffered Saline with added Tween20 (TBST) and 2% (w/v) skim milk or 3-5% (w/v) BSA protein for 15 minutes at room temperature, followed by incubation overnight at 4°C with a primary antibody (Table S3). Next, the membrane was incubated with species-specific horseradish peroxidase-conjugated immunoglobulin secondary antibody (Jackson Immuno Research or Agilent Technologies) for 1 hour at room temperature. Bands were visualised using Bio-Rad ChemiDocTM MP Imaging System and enhanced chemiluminescence (ECL+; Amersham Biosciences) or ImageQuant LAS 4000. Densitometric analysis was performed using Image LabTM Software, version 4.0 (Bio-Rad, USA; RRID: SCR_014210) or ImageQuant TL, version 8.1 software (GE Healthcare, USA; RRID: SCR_018374). Coomassie brilliant blue (18) or MEMCode reversible protein staining (20) were used as loading controls, and for each sample set, a representative membrane from the immunoblotting is shown.

### *In silico* analysis of skeletal muscle ageing time course

Gene expression data (GSE145480) ^56^ were extracted from SarcoAtlas (https://sarcoatlas.scicore.unibas.ch) to examine Rac1 transcript expression profile in the gastrocnemius muscle of 8, 18, 22, 24, 26 and 28 months-old male, C57BL/6JRj mice. Gene expression data was calculated as Log2(FC) relative to the 8-month samples.

### Analysis of Single Nucleotide Polymorphism (SNP) variants in FinnGen study

A human case-control study was conducted within the FinnGen study to assess the effects of SNP variants in Rac1. The FinnGen study is a large-scale genomics initiative that has analysed over 500,000 Finnish biobank samples and correlated genetic variation with health data to understand disease mechanisms and predispositions. The project is a collaboration between research organisations and biobanks within Finland and international industry partners. All participants in FinnGen provided informed consent. The study protocol and data collection have been previously described^57^.

Analyses were conducted in October 2025 using the FinnGen DF12 dataset (r12.finngen.fi). Associations between SNP variants in Rac1 and pre-selected phenotypes: “other disorders of carbohydrate metabolism”, “mixed hyperlipidaemia”, “hyperlipidaemia, other/unspecified”, “diseases of the myoneural junction and muscle”, and “muscle wasting and atrophy” were evaluated using REGINIE v.3.1.3. Logistic regression models were used with age, sex, the first 10 principal components of ancestry, FinnGen chip version and genotyping batch as covariates. Non-normally distributed traits were log-transformed before association. P-values were corrected for multiple comparisons using a Bonferroni p-value threshold of P < 0.05/5 (P < 0.01).

### Data visualisation

Data are presented as mean ± SEM with individual data points shown. Paired data points are connected with a straight line. Heatmaps were visualized using the R packages ComplexHeatmap (RRID:SCR_017270) ^58^ and circlize (RRID:SCR_002141) ^59^.

### Statistical Analyses

Outliers were defined as mean ±2SD. Excluded outliers are indicated in the figure legends. **S**tatistical tests varied according to the dataset being analysed, and the specific tests used are indicated in the figure legends. If a significant one-way ANOVA, Dunnett’s post hoc test was used to compare the experimental group to the control group. If a significant interaction was found in two-way repeated measures ANOVA or mixed model analysis, follow-up simple main effects were adjusted using Šídák’s confidence interval adjustment. Using the R package fgsea (RRID:SCR_020938) ^60^, the muscle proteomic data underwent gene set enrichment analysis (GSEA) using the log2 fold change between Rac1 imKO-Ctrl as the input ranked list and using MSigDB collections ‘H’ (hallmark gene sets) and ‘C2’ (curated gene sets) ^61–63^ and MitoCarta3.0 “MitoPathways” ^64^ to test for enrichment. P < 0.05 was considered statistically significant. 0.05 ≤ P < 0.1 was considered a tendency. Except for GSEA performed in R (RRID:SCR_001905), all statistical analyses were performed using IBM SPSS Statistics, version 30.0.0.0 (RRID:SCR_016479).

## Results

### Rac1 is implicated in muscle wasting with age in mice

To understand the implications of Rac1 in muscle wasting with age, we first determined Rac1 expression with ageing in mouse skeletal muscle. *In silico* analysis via the SarcoAtlas database (GEO accession number: GSE145480) ^56^ revealed increased Rac1 transcript level at 26 and 28 months of age (26 months: 15%; 28 months: 11%; Fig. 1A), compared with 8 months of age male C57BL/6JRj mice. The increase in Rac1 transcript level was concomitant with the mice reaching a stage of loss of muscle mass and strength mimicking sarcopenia ^56^. This upregulation of Rac1 transcript level seemed protective, as muscle depletion of Rac1 led to muscle wasting in middle-aged mice (Fig. 1B). This was evident as 12.7 ± 0.3 months old (mean ± SD) middle-aged muscle-specific inducible Rac1 knockout (Rac1 imKO) mice displayed lower muscle mass than littermate controls (Gastrocnemius: -10%; Quadriceps: -7%; Fig. 1B). Muscle mass was unaffected in 4.2 ± 0.6 months old adult Rac1 imKO mice (Fig. 1B) consistent with previous reports ^24^. The Rac1-dependent loss of muscle mass occurred before the age-related decrements in muscle mass and function set in ^56^, indicative of a Rac1-dependent mechanism of muscle wasting. Body weight, lean (internal organs, bone, and muscle), and fat mass were unaffected by the lack of skeletal muscle Rac1 protein in middle-aged mice (Fig. 1C). Intramyocellular protein E3 ubiquitin ligases previously ^65–67^ implicated in muscle wasting such as MuRF1 protein content (Fig. 1D, representative blots in E) and *Musa1* mRNA content (Fig. 1F), were unaffected by Rac1 deficiency (-85%; Fig. 1D) compared to control muscles. These findings implicate Rac1 deficiency in muscle mass regulation via unknown mechanisms not involving traditional muscle atrophy factors such as atrogenes.

**Figure 1.**
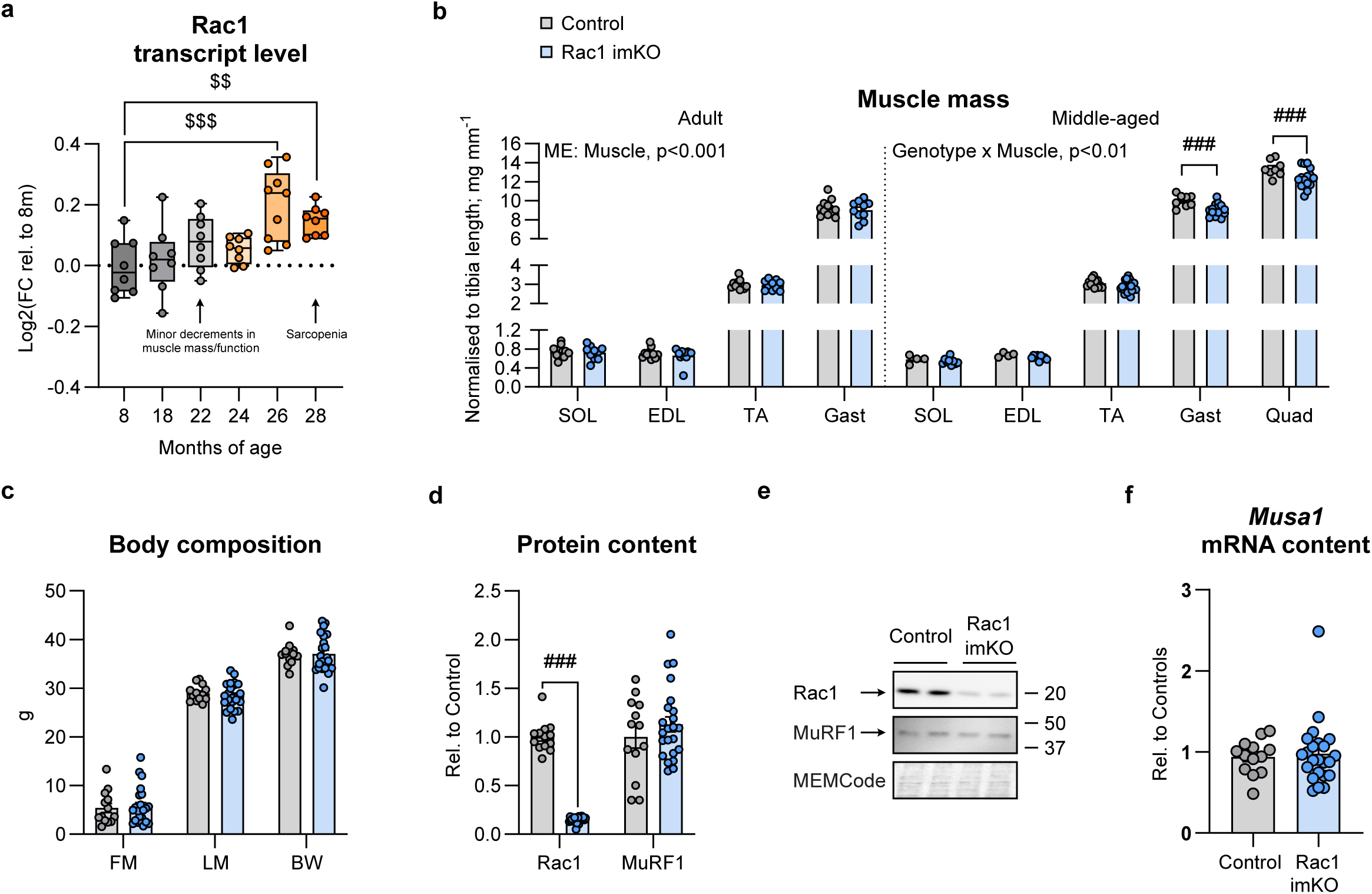
Rac1 is implicated in muscle wasting with age in mice. (A) Rac1 transcript level in the gastrocnemius muscle of male C57BL/6JRj mice at the indicated ages. Transcriptome previously published in the SarcoAtlas database (GEO accession number: GSE145480) ^56^. *n = 8/8/8/8/9/8* (8m/18m/22m/24m/26m/28m). For 28m, *n = 1* outlier excluded. Data evaluated with a one-way ANOVA. (B) Muscle mass normalised to tibia length in adult or middle-aged inducible muscle-specific Rac1 knockout (imKO) male mice or littermate controls. Adult Control, *n = 11/11/11/11* (SOL/EDL/TA/Gastrocnemius/Quadriceps); Adult Rac1 imKO, *n = 10/10/10/10*; Middle-aged Control, *n = 4/4/13/8/8*; Middle-aged Rac1 imKO, *n = 9/9/23/15/15*. Data were evaluated with a mixed model analysis for adult and middle-aged mice separately. (C) Body composition (FM: fat mass; LM: lean mass; BW: body weight) in middle-aged mice in gram. *n = 13/23* (Control/Rac1 imKO). (D) Rac1 and MuRF1 protein content. *n = 13/23* (Control/Rac1 imKO). (E) Representative blots showing (D) and control MEMCode protein staining. (F) *Musa1* mRNA content. If not otherwise indicated, data were evaluated with a Student’s t-test. Interactions are indicated in the panels. Significant interaction in mixed model analysis, significant one-way ANOVA or significant Student’s t-test: Effect of Rac1 imKO ### (p < 0.001); Effect of age $$/$$$ (p < 0.01/0.001). Data are presented as mean ± SEM with individual data points shown.

### Rac1 imKO muscles are enriched in proteins involved in fatty acid metabolism and oxidative phosphorylation

Having established that the deletion of Rac1 in mouse muscle led to muscle wasting with age, we next investigated the molecular mechanisms underlying this effect. To identify molecular alterations that precede muscle loss, we investigated 4.3 ± 1.1 months old adult Rac1 imKO mice. We first took an unbiased mass spectrometry approach to explore the proteomic changes in response to lack of muscle Rac1 protein in mice (Fig. 2A). Total protein abundance was obtained for 2311 proteins in five biological replicates (Fig. 2B). Expectedly, Rac1 was reduced in gastrocnemius muscle of Rac1 imKO mice compared to littermate controls (p < 0.05 adjusted for multiple testing and ±1.5-fold) and gene set enrichment analysis (GSEA) revealed depletion of gene sets associated with Rho GTPase signalling (Fig. S1A). Confirming Rac1’s involvement in cytoskeletal events, GSEA also revealed depletion of gene sets associated with pathways regulating the actin cytoskeleton, focal adhesions, and extracellular matrix regulation (Fig. 2C, Fig. S1A), for the first time cementing *in vitro* muscle cell data ^8^ on the role of Rac1 on actin cytoskeleton regulation in mature skeletal muscle. Gene sets associated with myogenesis were depleted (Fig. 2D), indicating that muscle-building processes were impaired. Interestingly, GSEA also showed that gene sets associated with fatty acid metabolism and oxidative phosphorylation were enriched at the proteome level among Rac1-deficient muscles compared to controls (Fig. 2C-D, Fig. S1A), exposing potential new roles for Rac1 in skeletal muscle biology.

**Figure 2:**
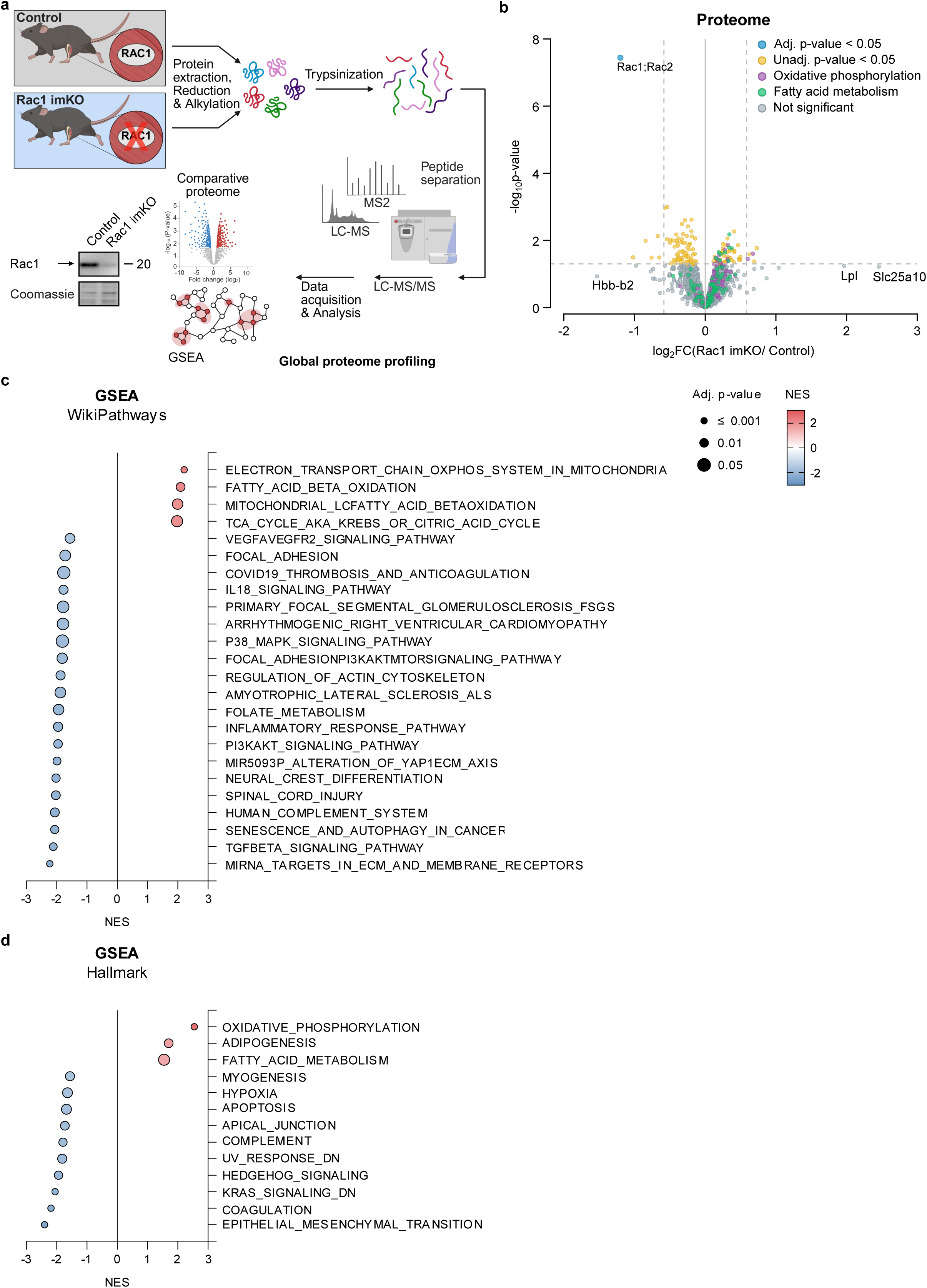
Rac1 imKO muscles are enriched in proteins involved in fatty acid metabolism and oxidative phosphorylation. (A) Schematic of the proteome sample preparation workflow. (B) Volcano plot of all protein groups detected with quantitative MS-based proteomics in gastrocnemius muscle from adult inducible muscle-specific Rac1 knockout (imKO) mice or littermate controls. *n = 4/1* (male/female). Unadjusted p-values. 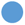 Adjusted p-values < 0.05, 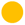 Unadjusted p-value < 0.05, 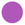 Hallmark Oxidative phosphorylation gene set, 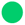 Hallmark Fatty acid metabolism gene set, 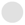 Not significant. (C) WikiPathways Gene Set Enrichment Analysis (GSEA) and (D) Hallmark GSEA showing Normalised Enrichment Score (NES) values for each statistically significant gene set and adjusted p-values.

Collectively, our exploration indicates that muscle Rac1 orchestrates proteomic alterations associated not only with anticipated cytoskeletal dynamics but also, unexpectedly, with the modulation of proteins governing fatty acid metabolism and oxidative phosphorylation, potentially involving Rac1 in mitochondrial dysfunction ^6^, fat infiltration and alterations in fatty acid metabolism ^68–70^, hallmarks of ageing and sarcopenia.

### Rac1 imKO mice demonstrate increased mitochondrial content and impaired lipid handling and fatty acid oxidative capacity in muscle

The discovery that proteins involved in fatty acid metabolism and oxidative phosphorylation were upregulated in Rac1 imKO muscles prompted us to examine potential mitochondrial changes and explore Rac1’s hitherto unknown role in fatty acid oxidation. Using immunoblotting, we confirmed a marked upregulation of subunits of the OXPHOS complex in Rac1-deficient quadriceps muscle compared to control (CI subunit NDUFB8: +149%, CII subunit SDHB: +169%, CIII subunit UQCRC2: +455%, CIV subunit MTCO1: +92%, CV subunit ATP5A: +50%; Fig. 3A, representative blots in 3B). We also observed alterations in mitochondrial morphology of Rac1 imKO muscles compared to controls using transmission electron microscopy. In both the subsarcolemmal (SS, Fig. 3C) and intermyofibrillar (IMF; Fig. 3D) areas, several of the mitochondria in the Rac1 imKO muscles appeared enlarged and damaged, which was not apparent in any of the analysed control muscles. When quantified, these morphological changes in Rac1-deficient muscles manifested as a +467% increase in SS (Fig. 3E) and +166% increase in IMF (Fig. 3F) mitochondrial volume densities. The increased mitochondrial volume was also indicated by a highly elevated surface area of the cristae membrane per muscle volume (+250%, Fig. 3G). Rac1-deficiency caused a similar directional change, however SS mitochondrial volume increased primarily due to an increase in the number of mitochondria (+253%; Fig. 3H) with the mean mitochondrial area statistically unchanged (Fig. 3I), while IMF mitochondria showed no change in number (Fig. 3J), but with increased mean mitochondrial area (+83%; Fig. 3K) in Rac1 imKO muscle compared to controls. Although Rac1 deficiency induced similar directional changes in SS and IMF mitochondria (Fig. S2A–H), morphological alterations were more pronounced in IMF mitochondria, with increased perimeter (+30%, p = 0.063; Fig. S2E), inner diameter (+28%, p = 0.075; Fig. S2G) and maximal Feret diameter (+30%, p = 0.052; Fig. S2H) versus controls, consistent with their larger area. Elevated % damaged mitochondria was observed in 3 out of 5 of the Rac1 imKO muscles, compared to control muscles (Fig. S2I). This altogether suggests that Rac1 deficiency leads to enlarged or increased number of mitochondria, depending on the mitochondrial subfraction, and to some extent, damaged mitochondria.

**Figure 3:**
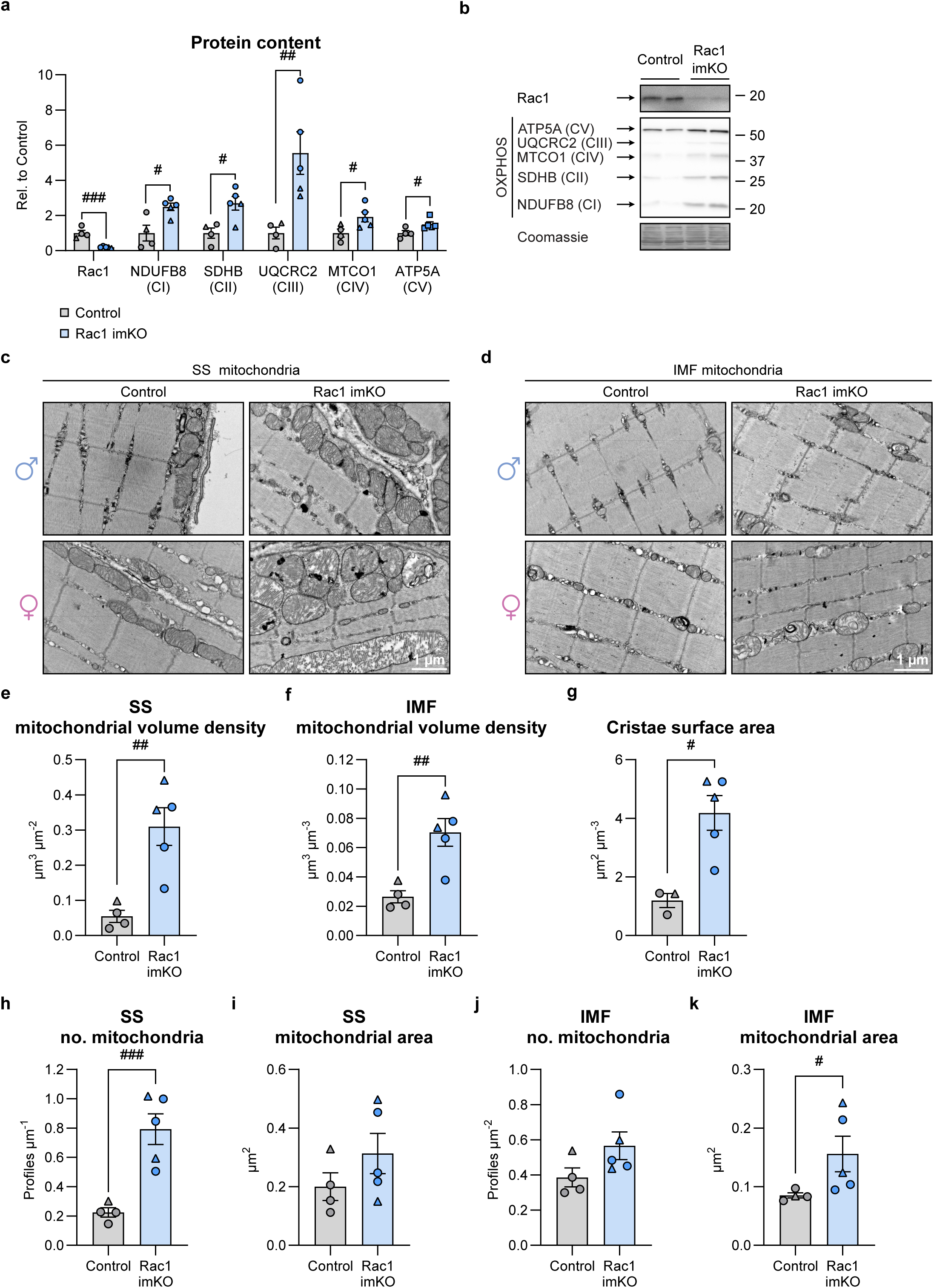
Rac1 imKO mice demonstrate increased mitochondrial content in muscle. (A) OXPHOS (subunits of mitochondrial oxidative phosphorylation complexes [CI: NDUFB8; CII: SDHB, CIII: UQCRC2; CIV: MTCO1, CV: ATP5A]) protein content in quadriceps muscle from adult inducible muscle-specific Rac1 knockout (imKO) mice or controls. Control, *n = 3/1* (male/female); Rac1 imKO, *n = 3/2*. (B) Representative blots showing (A), Rac1 protein content and control Coomassie staining. (C-D) Representative Transmission Electron Microscopy (TEM) images showing skeletal muscle mitochondria in quadriceps muscle in the (C) subsarcolemmal (SS) and (D) intermyofibrillar (IMF) areas in male and female mice. Scale bar, 1 µm. (E) SS and (F) IMF mitochondrial volume density. (G) Mean surface area of cristae membrane per volume of fibre. Data were not obtained for *n = 1* male control mouse. (H) SS total mitochondrial number and (I) and mean mitochondrial area. (J) IMF total mitochondrial number and (K) and mean mitochondrial area. Data were evaluated with a Student’s t-test: Effect of Rac1 imKO #/##/### (p < 0.05/0.01/0.001). Data are presented as mean ± SEM with individual data points shown. ◯ Male, △ Female.

Mitochondria are, along with peroxisomes, the major sites for fatty acid β-oxidation. To understand the functional consequences of these mitochondrial alterations, we investigated the muscle metabolic profile. Loss of Rac1 caused some re-distribution of lipids within skeletal muscle, including diacylglycerol (DAG), TG, acylcarnitine, sphingomyelin, and ceramide species (cytosolic 1,3-DAG(16:0/18:2): +103%, p = 0.081; membrane-associated 1,3-DAG(16:0/20:4): - 63%; membrane-associated 1,3-DAG(18:0/18:1): -63%, p =0.098; acylcarnitine(14:0): +73%, p = 0.068; Fig. S3A-H). Next, as exercise robustly mobilises metabolic turnover, including fatty acid oxidation and lipid uptake in muscle ^71^, we studied the role of Rac1 in these processes. In response to treadmill running, a total of 14 metabolites (out of 483 metabolites) were differentially regulated in the gastrocnemius muscle of Rac1 imKO mice compared to controls (Fig. S4A). Rac1 deficiency caused 11 out 13 acylcarnitine metabolites to accumulate, including a tendency to upregulation of acetylcarnitine (+9%, p = 0.081; Fig. S4B), as well as high levels of the monohydroxy fatty acids 2-hydroxydecanoate (+26%; Fig. S4C), 3-hydroxyoctanoate (+29%; Fig. S4C), and 3-hydroxypalmitate (+53%; Fig. S4C), which could reflect that mitochondrial β-oxidation and/or fatty acid import to the mitochondria were not occurring to a sufficient extent, resulting in an accumulation of unoxidized carnitine conjugated fatty acids and monohydroxy fatty acids. A limited carbon flux to the TCA cycle from fatty acids could result in the consequent depletion of the fatty acid and branched-chain amino acid (BCAA) metabolites methylmalonate (MMA; -51%; Fig. S4B), propionylcarnitine (-34%; Fig. S4B), isovalerylcarnitine (-31%, p = 0.057; Fig. S4D), and beta-hydroxyisovalerate (-40%; Fig. S4D). The levels of detected medium chain fatty acids, long chain fatty acids, branched chain fatty acids and polyunsaturated fatty acids were similar (Fig. S4E), suggesting no difference in fatty acid availability intramuscularly between Rac1-deficient muscle and control.

Consistent with impaired lipid handling, contraction-stimulated palmitate oxidation was blunted in Rac1 imKO mice, representing a 62% lower increase compared to controls (ΔPalmitate oxidation; Fig. 4A). The contraction-induced increase in palmitate uptake was also lower in Rac1 imKO muscles than in controls (ΔPalmitate uptake: -61%; Fig. 4B). Beyond lipid uptake, contraction also mobilises intramyocellular triacylglycerol (TG) for oxidation and we therefore assessed TG utilisation during *in-situ* contraction. The contraction-induced decrease in muscle TG was diminished in muscle deficient of Rac1 (ΔMuscle TG: -102%, Fig. 4C), altogether indicating a compromised lipid handling and oxidative capacity in Rac1 imKO muscle. *In situ* contraction-induced glycogen breakdown in the quadriceps muscle was similar between groups (Fig. 4D) in alignment with previous studies ^27^. This suggests a role for Rac1 in fatty acid breakdown and the increase in fatty acid oxidation in response to muscle contraction. Taken together, these data implicate Rac1 in skeletal muscle mitochondrial alterations of morphology and changes in fatty acid metabolism, both of which precede muscle mass loss in Rac1-deficient muscle.

**Figure 4:**
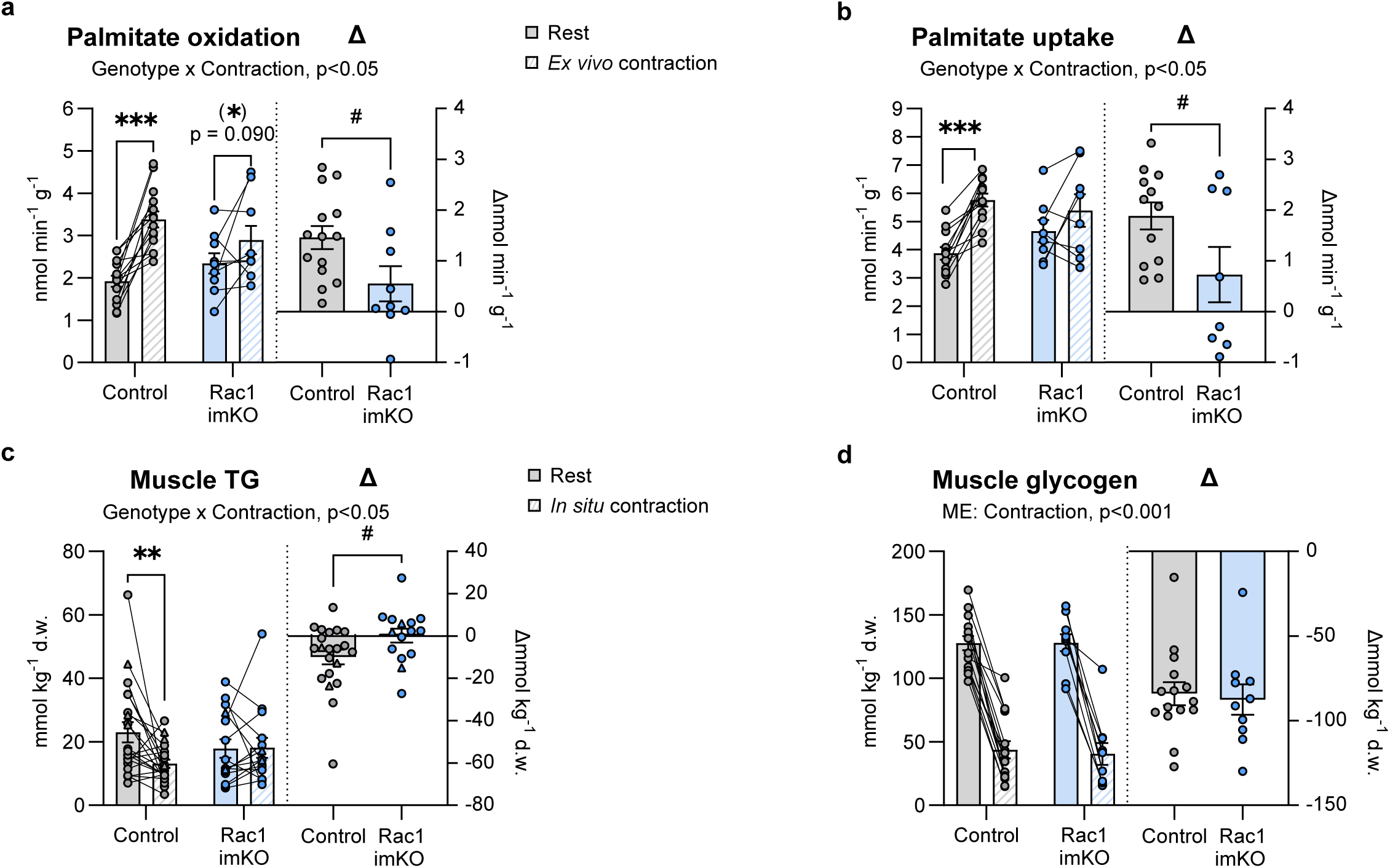
Rac1 imKO mice demonstrate impaired lipid handling and fatty acid oxidative capacity in muscle. (A) Contraction-stimulated (600 msec/3.3 sec, 30 Hz, 25 min) palmitate oxidation and (B) palmitate uptake in isolated soleus muscle from adult male inducible muscle-specific Rac1 knockout (imKO) mice or littermate controls. Control, *n = 14/14* (Rest/Contraction); Rac1 imKO, *n = 9/9*. For Control, n = 1 outlier excluded. Data for palmitate uptake were not obtained for *n = 2* control mice and *n = 1* Rac1 imKO mouse. (C) *In situ* contraction-stimulated (600 msec/3.3 sec, 30 Hz, 25 min) muscle triacylglycerol (TG) and (D) glycogen in quadriceps muscle from Rac1 imKO mice or littermate controls. d.w., dry weight. Muscle TG: Control, *n = 21/21* (Rest/Contraction; 18/3 Male/female); Rac1 imKO, *n = 15/15* (12/3). Glycogen (male only): Control, *n = 15/15* (Rest/Contraction); Rac1 imKO, *n = 10/10*. Contraction-stimulated data were evaluated with a two-way repeated-measures (RM) ANOVA. Delta values were evaluated with a Student’s t-test. Main effects and interactions are indicated in the panels. Significant interaction in two-way RM ANOVA and significant Student’s t-test: Effect of contraction (*)/**/*** (p < 0.1/0.05/0.001); Effect of Rac1 imKO # (p < 0.05). Data are presented as mean ± SEM with individual data points shown. Paired data points are connected with a straight line. ◯ Male, △ Female.

### The lack of muscle Rac1 lowers mitochondrial respiratory capacity in middle-aged mice

Intrigued by a potentially novel role for Rac1 in mitochondrial volume and fatty acid oxidation, we wanted to dive deeper into Rac1’s potential mechanistic role in regulating mitochondrial respiration. Inquisitive by the recent discovery of mitochondria-localised Rac1 with Bcl-2 in the mitochondria in neuronal cells ^72^, we determined this in mature skeletal muscle. We found that the majority of Rac1 localised to the cytosol with a minor proportion in the mitochondria in unstimulated skeletal muscle (Fig. 5A, representative blot in 5B). Overall, 15% ± 2 (mean ± SEM) of the Rac1 protein in the muscle localised to the mitochondria (Fig. 5A). Given that mitochondrial energetics associate with muscle quality ^6^, we wanted to know if the lack of skeletal muscle Rac1 would accelerate a decline in mitochondrial respiration, potentially explaining the muscle wasting we observed in the middle-aged Rac1 imKO mice. To this end, we studied the middle-aged Rac1 imKO mice again at a time when mitochondrial respiratory flux is already starting to decline in muscle ^73^. High-resolution respirometry in permeabilized gastrocnemius skeletal muscle fibre bundles revealed a reduced mitochondrial respiratory capacity in the Rac1 imKO muscles compared to controls (Fig. 5C). This was evident when adding malate and pyruvate to assess LEAK respiration (-32%), ADP (5 mM) to assess state 3 respiration (-25%, p = 0.059), glutamate to assess complex I linked respiratory activity (-28%, p = 0.055) and succinate to assess complex I + II linked respiratory capacity (-27%). Citrate synthase (CS) activity, a marker of mitochondrial content ^74,75^, was increased in middle-aged Rac1 imKO muscles compared to controls (+23%; Fig. 5D), supporting increased mitochondrial content. Normalising to CS activity, muscle respiratory capacity was further decreased per mitochondrial content in Rac1-deficient mice relative to control mice (Mal + Pyr: -46%; ADP 0.25 mM: -33%; ADP 5.0 mM: -40%; Glut: -42%; Succ: -42%; AMA: -48%; Fig. 5E). In alignment of increased CS activity, and in further support of increased mitochondrial content, protein content of the CV subunit ATP5A of the OXPHOS complex was upregulated in gastrocnemius muscle of the middle-aged Rac1 imKO mice compared to control mice (+28%; Fig. 5F, representative blot in G), similar to the younger adult mice.

**Figure 5:**
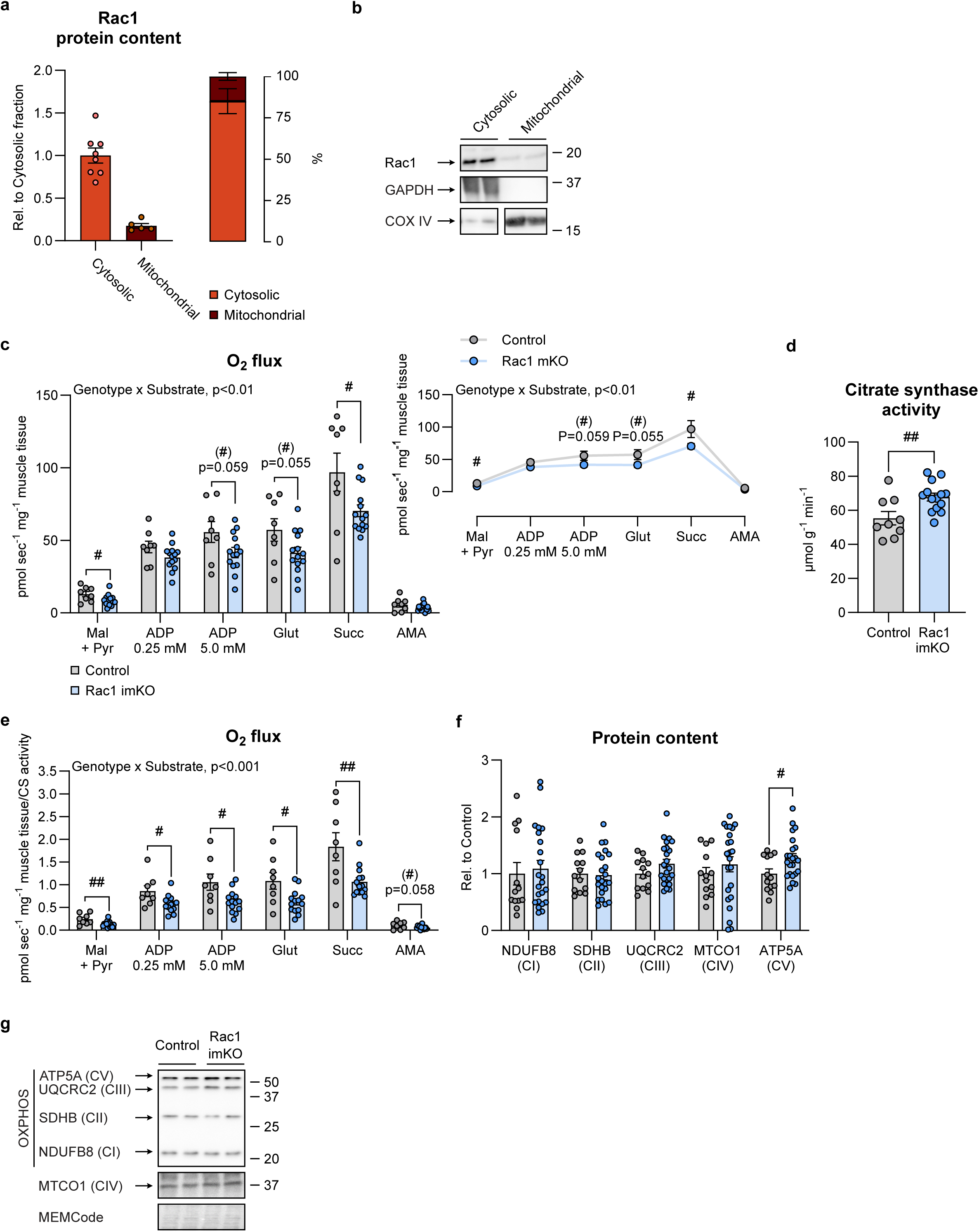
The lack of muscle Rac1 lowers mitochondrial respiratory capacity in middle-aged mice.(A) Rac1 protein content in the cytosolic and mitochondrial fractions, isolated from female C57BL/6N quadriceps mouse muscle. *n = 8/5* (Cytosolic/Mitochondrial). (B) Representative blot showing (A) including GAPDH (marker of cytosolic fraction) and COX IV (marker of mitochondrial fraction). The representative blots showing COX IV are cropped from the same membrane. (C) High-resolution respirometry in permeabilised gastrocnemius muscle fibers. *n = 8/14* (Control/Rac1 imKO). For Control, n = 1 outlier excluded. Mal: Malate, Pyr: Pyruvate, Glut: Glutamate, Succ: Succinate, AMA: Antimycin A. (D) Citrate synthase (CS) activity. *n = 9/14* (Control/Rac1 imKO). (E) Respiratory capacity in (C) related to CS activity in (D). (F) OXPHOS (subunits of mitochondrial oxidative phosphorylation complexes [CI: NDUFB8; CII: SDHB, CIII: UQCRC2; CIV: MTCO1, CV: ATP5A]) protein content. *n = 13/23* (Control/Rac1 imKO). (G) Representative blots showing (F) and control MEMCode staining. O_2_ flux data were evaluated with a two-way repeated-measures (RM) ANOVA. Protein content and CS activity were evaluated with a Student’s t-test. Interactions are indicated in the panels. Significant interaction in two-way RM ANOVA or significant Student’s t-test: Effect of Rac1 imKO (#)/#/## (p < 0.1/0.05/0.01). Data are presented as mean ± SEM with individual data points shown. Paired data points are connected with a straight line.

Having identified mitochondrial alterations and changed fatty acid metabolism preceding muscle loss in Rac1-deficient muscle, we wanted to dig deeper into the molecular mechanisms for the accelerated decline in respiratory capacity in middle-aged Rac1 imKO mice. Total protein abundance was obtained for 4645 proteins in four biological replicates in proteomic analysis of gastrocnemius muscle from these older middle-aged mice (Fig. 6A). Similar to the younger adult mice, GSEA showed that gene sets associated with fatty acid metabolism and oxidative phosphorylation were enriched at the proteome level among the muscles from the older middle-aged Rac1 imKO mice compared to controls (Fig. 6B-C, Fig. S5A). These data underscore the mitochondrial and fatty acid metabolic phenotype in muscle deficient of Rac1, further strengthened by an overlap of 30 enriched or depleted gene sets between adult and middle-aged Rac1 imKO mice (Fig. S5B). Yet, in the older middle-aged mice, Rac1 deficiency was also associated with enrichment of mitochondrial complex I assembly and disorders of mitochondrial homeostasis dynamics protein import and quality control (Fig. 6C), and enrichment of mitochondrial protein degradation, complex I biogenesis, complex III assembly, and maturation of TCA enzymes and regulation of TCA cycle (Fig. S5A). Further, enrichment analysis based on the MitoPathways from the MitoCarta3.0 database ^64^, revealed enrichment of the gene set associated with mt-tRNA syntheases (Fig. 6D) in middle-aged Rac1 imKO muscle. Together, these data suggest increased turnover of mitochondrial proteins, indicating a worsened mitochondrial phenotype with age in Rac1 imKO muscle consistent with impaired mitochondrial respiratory capacity (Fig. 5C) and muscle wasting (Fig. 1D) observed in middle-aged mice. We next measured the content of proteins involved in mitochondrial fission (DRP1), fusion (OPA1, MFN2), mitophagy (BNIP3, PINK1, PARKIN), and autophagy (ULK1, pULK1 S555 and pULK1 S757, p62, p-p62 T269/S272) (Fig. 6E, representative blots in F). We did not observe any major changes in protein content, apart from a slight upregulation of OPA1 (+15%, p = 0.090) and PINK1 (+15%) protein content in the Rac1 imKO muscle. We observed an increase in pULK1 S757 ULK (mTORC1 site; +18%) but not pULK1 S555 (AMPK site), indicative of decreased autophagy. The most pronounced effect was observed on p-p62 T269/S272, which was 42% lower in the Rac1 imKO muscle than in control mice, indicative of reduced mTORC1 activity ^76^, collectively suggesting repressed cell growth and dysfunctional autophagy/mitophagy. Altogether, these findings indicate that mitochondrial oxidative capacity is decreased in middle-aged Rac1 imKO gastrocnemius muscle, not due to fission-fusion defects, but rather possibly driven by altered mitophagy and autophagy regulation, which could lead to accumulation of damaged and dysfunctional mitochondria.

**Figure 6:**
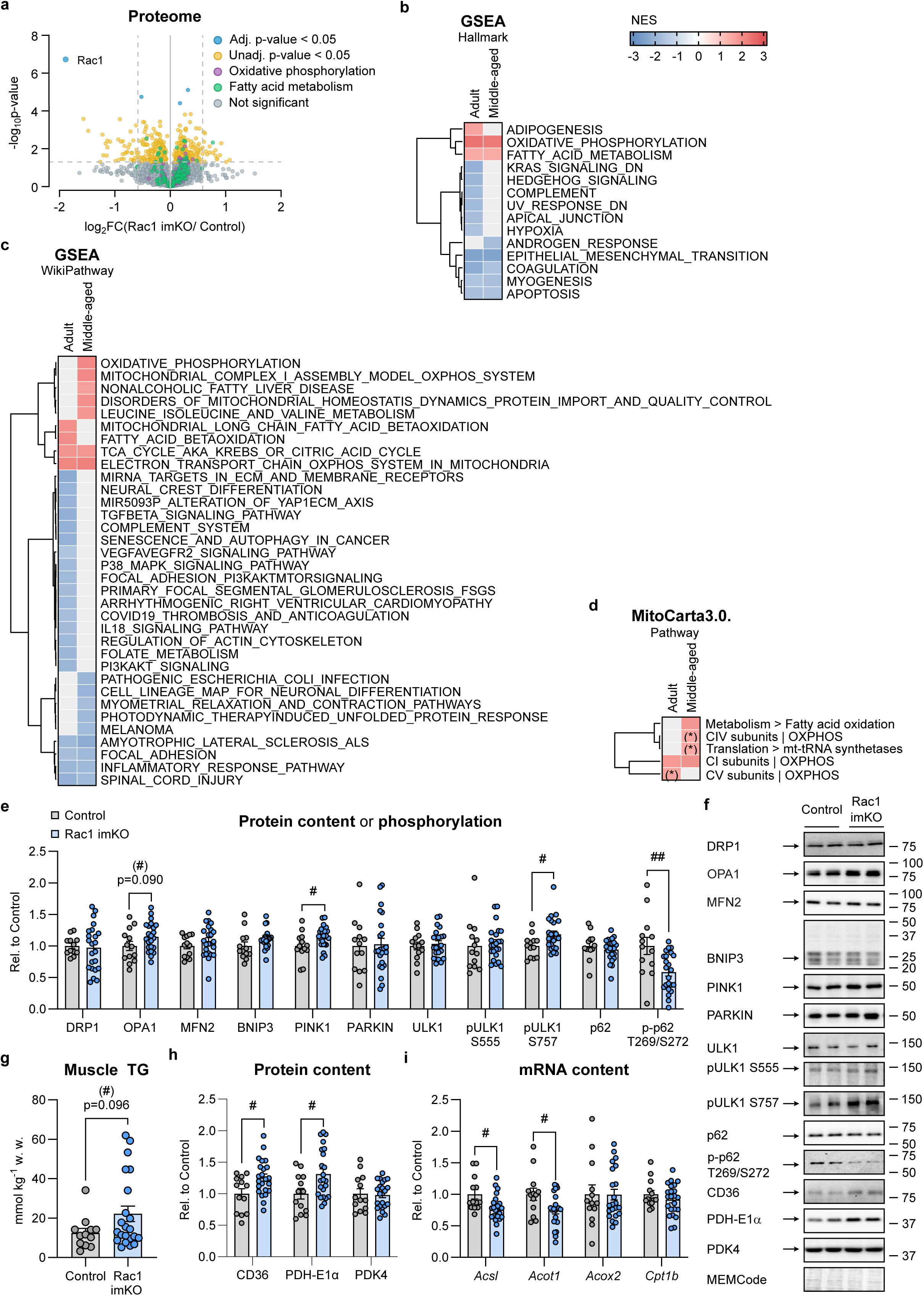
Rac1 deficiency is associated with molecular changes in mitophagy, autophagy and fatty acid metabolism signalling. (A) Volcano plot of all protein groups detected with quantitative MS-based proteomics in gastrocnemius muscle from middle-aged male inducible muscle-specific Rac1 knockout (imKO) mice or littermate controls. *n = 4/4* (Control/Rac1 imKO). Unadjusted p-values. 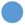 Adjusted p-values < 0.05, 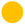 Unadjusted p-value < 0.05, 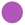 Hallmark Oxidative phosphorylation gene set, 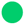 Hallmark Fatty acid metabolism gene set, 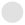 Not significant. (B) Hallmark Gene Set Enrichment Analysis (GSEA), (C) WikiPathways GSEA, and (D) MitoCarta3.0 MitoPathway GSEA showing Normalised Enrichment Score (NES) values for each statistically significant gene set in adult or middle-aged gastrocnemius mouse muscle. (E) DRP1, OPA1, MFN2, BNIP3, PINK1, PARKIN, ULK1, and p62 protein content and phosphorylated (p)ULK1 S555, pULK1 S757 and p-p62 T269/S272. *n = 13/23* (Control/Rac1 imKO). For DRP1, data were not obtained for *n = 1* control mouse. (F) Representative blots showing (E+H) and control MEMCode protein staining. (G) Muscle triacylglycerol (TG) in gastrocnemius muscle. w.w., wet weight. For controls, *n = 1* outlier excluded. For Rac1 imKO, data were not obtained for *n = 1*. (H) CD36, PDH-E1α, and PDK4 protein content. (I) *Acsl, Acot1, Acox2,* and *Cpt1b* mRNA content. For *Acox2*, *n = 1* outlier excluded in Rac imKO. Data were evaluated with a Student’s t-test. Significant Student’s t-test: Effect of Rac1 imKO (#)/#/## (p < 0.1/0.05/0.01). Data are presented as mean ± SEM with individual data points shown.

The lower mitochondrial respiration in middle-aged Rac1 imKO muscle compared to control aligned with reduced activation of fatty acid oxidation and a lower usage of muscle TG in response to muscle contraction in the adult Rac1 imKO mice. Reduced mitochondrial respiration and lower fatty acid oxidation could, over time, result in the accumulation of muscular TG. Accordingly, middle-aged Rac1 imKO muscle tended to have increased muscle TG (+78%, p = 0.096; Fig. 6G), as well as an upregulation of fatty acid transporter, CD36 protein content (+25%; Fig. 6H). Moreover, we observed down-regulated acyl-CoA synthetase long-chain family member 1, *Acsl1* (-22%) and acyl-CoA thioesterase 1, *Acot1* (-25%) mRNA content in Rac1-deficient muscle, while *Acox2* and *Cpt1b* mRNA content was unaffected (Fig. 6I), altogether implicating Rac1 in lipid storage, uptake, and oxidation mechanisms within the muscle tissue. Also, Rac1-deficiency increased PDH-E1α (+31%) but not PDK4 protein content compared to control muscles (Fig. 6H). The PDH complex is the rate-limiting multienzyme complex responsible for irreversible decarboxylation of pyruvate acetyl-CoA, thereby linking glycolysis to the citric acid cycle, thus the upregulation could potentially compensate for the impaired mitochondrial substrate regulation. Taken together, these data for the first time implicate Rac1 in muscle lipid handling, ultimately causing muscle TG accumulation, which could possibly be involved in the muscle wasting observed in middle-aged Rac1-deficient mice.

### Rac1 associates with sarcopenia-driven alterations in human skeletal muscle as well as metabolic and muscle-wasting diseases

Having established that Rac1 deficiency caused skeletal muscle wasting, mitochondrial dysfunction and alterations in fatty acid metabolism in mice, we next examined human muscle. Underlining the clinical importance of our mouse findings, we observed that Rac1 protein content was increased in muscle biopsy samples obtained from old, sarcopenic subjects compared to young subjects (+41%; Fig. 7A, representative blot in 7B). It has been established that sarcopenia is caused primarily by type II fibre atrophy ^77–80^. When combining young and old, sarcopenic subjects, we observed that Rac1 protein content was negatively correlated with quadriceps CSA (p < 0.05, r = -0.475; Fig. 7C) and type II fibre CSA (p = 0.051, r = -0.466; Fig. 7D), potentially implicating Rac1 in sarcopenia-driven alterations in human skeletal muscle. Assessing mitochondrial OXPHOS protein content and stratifying for age, we found that in non-sarcopenic young subjects, Rac1 protein content correlated positively with CIII subunit UQCRC2 protein content (p < 0.05, r = 0.734; Fig. 7F) and tended to correlate negatively with CIV subunit MTCO1 protein content (p = 0.061, r = - 0.610; Fig. 7G). In old, sarcopenic patients Rac1 protein content correlated negatively with CI subunit NDUFB8 protein content (p < 0.05, r = -0.690; Fig. 7E), CIV subunit MTCO1 protein content (p < 0.001, r = -0.938; Fig. 7G) and CV subunit ATP5A protein content (p < 0.05, r = -0.704; Fig. 7H), associating Rac1 with protein content of mitochondrial respiratory complexes in human skeletal muscle. No correlations were found between Rac1 protein content and type I fibre CSA and OXPHOS CII subunit SDHB protein content (Fig. S6A-B). Finally, we also conducted a human case-control study within the FinnGen dataset containing genetic and phenotypic data of >500,000 individuals with a median age of 53 years, to assess the effects of Rac1 single-nucleotide polymorphism (SNP) variants. We found significant associations between Rac1 SNP variants and “other disorders of carbohydrate metabolism” (OR: 18.36, p = 1.80 × 10^-3^), “hyperlipidaemia, other/unspecified” (OR: 1.27, p = 1.50 × 10^-3^), “mixed hyperlipidemia” (OR: 2.92, p = 3.00 × 10^-4^) and “muscle wasting and atrophy” (OR: 4.44, p = 1.00 × 10^-5^) with higher probability of being diseased (Fig. 7I). On the contrary, a lower probability of disease was found between Rac1 SNP variants and “diseases of the myoneural junction and muscle” (OR: 0.89, p = 5.50 × 10^-4^; Fig. 7I). This human case-control study points towards a hitherto unrecognised role for Rac1 in lipid metabolism and muscle wasting diseases in humans. Altogether, these results indicate that Rac1 is implicated in sarcopenia-driven alterations in human skeletal muscle and is associated with metabolic and muscle-wasting diseases, providing important translational data for the role of Rac1 in age-related muscle decline and mitochondrial function (Fig. 8A).

**Figure 7:**
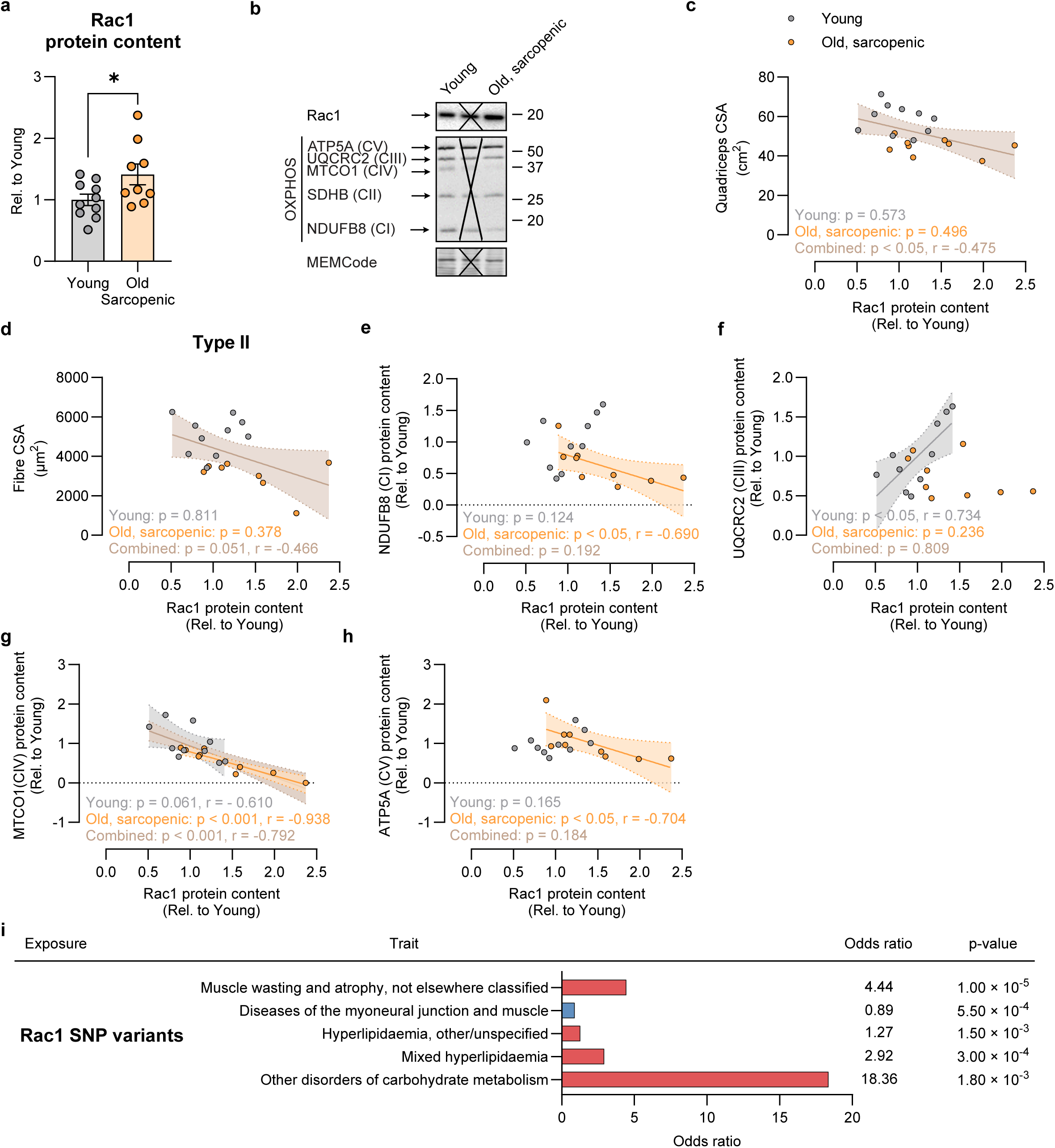
Rac1 associates with sarcopenia-driven alterations in human skeletal muscle as well as metabolic and muscle-wasting diseases. (A) Rac1 protein content in old, sarcopenic vastus lateralis muscle compared to young control. *n = 10/9* (Young/Old, sarcopenic). Data were evaluated with a Student’s t-test. (B) Representative blots showing (A) and control Coomassie staining. X, indicate lane not included in the sample set. (C) Correlation between Rac1 protein content and quadriceps cross-sectional area (CSA; quadriceps CSA was previously published ^28,29^)), (D) type II fibre CSA, (E; type II fibre CSA was previously published ^28,30^),) OXPHOS CI subunit NDUFB8 protein content, (F) CIII subunit UQCRC2 protein content, (G) CIV subunit MTCO1 protein content, and (H) CV subunit ATP5A protein content. *n = 10/9* (Young/Old, sarcopenic). For type II CSA, data were not obtained for *n = 1*. Data were evaluated with Pearson’s correlation. (I) Association between Rac1 SNP variants and indicated traits in FinnGen study participants calculated from linear regression models of association. Significant Student’s t-test: Old, sarcopenic vs. Young * (p < 0.05). Data are presented as mean ± SEM with individual data points shown.

**Figure 8:**
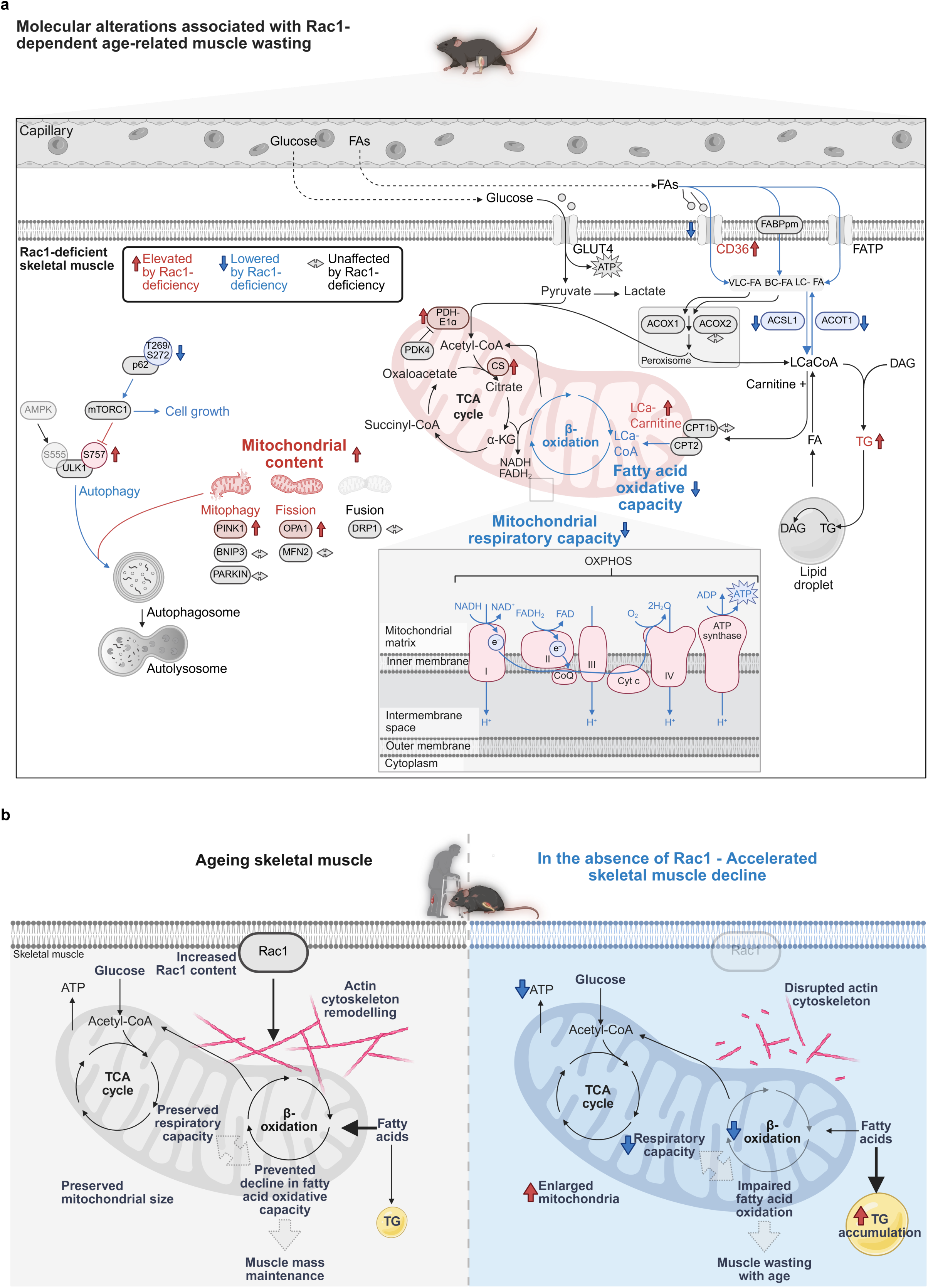
(A) Schematics of molecular alterations associated with Rac1-dependent age-related muscle wasting. (B) Working model in which muscle Rac1 content increases with age to protect against age-related muscle decline through a hierarchical cascade: Rac1-dependent actin remodelling preserves mitochondrial size and respiratory capacity. The protection of mitochondrial function prevents a decline in fatty acid oxidative capacity, ultimately contributing to muscle mass maintenance.

## Discussion

Our results shed new light on the role of Rac1 in skeletal muscle maintenance. We demonstrate that skeletal muscle Rac1 protein content increases with age in humans, likely serving as a compensatory mechanism against sarcopenia. Supporting this, muscle-specific Rac1 deficiency in mice impairs mitochondrial respiration and fatty acid metabolism, driving accelerated muscle wasting. Our findings support a model in which Rac1 counteracts sarcopenia through a hierarchical axis: Rac1-dependent actin remodelling preserves mitochondrial morphology and respiratory capacity, thereby maintaining fatty acid oxidative capacity and ultimately sustaining muscle mass (Fig. 8B). Five major new findings are derived from the present investigation: First, Rac1-deficient middle-aged mice displayed premature muscle wasting. Second, proteomic profiling confirmed that Rac1 loss disrupts cytoskeletal regulation while unexpectedly enriching pathways for fatty acid metabolism and oxidative phosphorylation, potentially as an adaptive response to mitochondrial positioning defects. Third, muscle-specific Rac1 deletion induced mitochondrial accumulation and morphological defects characteristic of impaired dynamics. These disruptions likely underpin the respiratory impairments observed in middle-aged mice, contributing to muscle wasting. Fourth, metabolomic and functional analyses demonstrated altered lipid profile, impaired lipid handling and reduced contraction-stimulated fatty acid oxidation and uptake despite elevated mitochondrial protein abundance. Finally, in sarcopenic human muscle, Rac1 protein content was elevated and negatively correlated with multiple mitochondrial respiratory complexes, while Rac1 SNP variants were associated with lipid metabolic and muscle-wasting diseases, underscoring translational relevance. Collectively, these findings position Rac1 as a previously unrecognised factor in the regulation of mitochondrial integrity and fatty acid metabolism in skeletal muscle with implications for muscle mass and thereby the pathophysiology of muscle decline and sarcopenia.

We demonstrate that Rac1 deficiency drives muscle wasting with age, supported by the depletion of myogenic gene sets, indicating an impairment of essential muscle-building processes. This adds to a growing body of evidence supporting a role for Rac1 in muscle mass regulation. In cardiac muscle, Rac1 is critical in the regulation of cardiac hypertrophy ^81–83^. In alignment, mice with decreased catalytic activity of Rho GTPase activator, Vav2, exhibit reduced skeletal muscle mass ^84^, and we recently documented impaired exercise training-induced hypertrophy in Rac1 imKO skeletal muscle ^24^. Moreover, age-related muscle wasting in skeletal muscle deficient of Rac1 downstream targets, PAK1/2 ^13^ combined with our finding of Rac1-dependent loss of muscle mass with age, implies a role for Rac1 in the protection against sarcopenia.

Rac1 deletion remodelled the muscle proteome beyond cytoskeletal and muscle-building pathways, enriching proteins involved in fatty acid metabolism and oxidative phosphorylation and revealing an unrecognised role for Rac1 in fatty acid metabolism. While the importance of Rac1 in the translocation of the glucose transporter, GLUT4, in skeletal muscle is well-established ^7,9–11,25–27,85,86^, the function of Rac1 in metabolic rewiring in skeletal muscle represents a previously unrecognised dimension of Rac1 signalling. In cancer cells, Rac1 reprograms glucose metabolism by promoting the transcription of glycolytic enzymes ^87,88^. Moreover, Rac1-dependent metabolic reprogramming is potentially achieved by coupling cytoskeletal dynamics to metabolic pathways. In endothelial MCF10A cells, PI3K-dependent activation of Rac1 leads to cytoskeletal reorganisation-dependent aldolase A activation, thereby determining glycolytic flux ^89^. While we did not determine Rac1-dependent regulation of glycolysis in skeletal muscle, our data points towards a role for Rac1 in fatty acid oxidation.

Our findings further identify a role for muscle Rac1 in the control of mitochondrial morphology and, in middle-aged mice, respiratory capacity. To the best of our knowledge, those results provide the first link between Rac1 and mitochondria in skeletal muscles, but it corroborates findings in L6-GLUT4myc myotubes, where siRNA-mediated depletion of Rac1 downstream effector PAK1 decreased respiration and mitochondrial copy number ^90^. The implication of Rac1 in mitochondrial homeostasis in skeletal muscle could be due to the known role of Rac1 in actin cytoskeleton organisation. Emerging studies suggest that the actin cytoskeleton is important for processes regulating mitochondrial morphology. For example, in U2OS cell, an actin-dependent step in mitochondrial fission was found to be mediated by the endoplasmic reticulum (ER)-associated formin inf2 ^91^, a protein that accelerates both actin polymerisation and depolymerisation. Thus, Rac1 may play a role in the communication between the SR and mitochondria via the actin cytoskeleton. Moreover, pharmacological inhibition of the Rac1 effector Arp2/3 disrupted actin polymerisation required for mitochondrial fusion in human fibroblasts and MDA-MB-231 cells ^92^, while shRNA-mediated knockdown of cortactin, cofilin, or Arp2/3 induced aberrant mitochondrial elongation in MEFs ^93^. Further corroborating an important implication of the actin cytoskeleton in mitochondrial regulation, the actin depolymerising agent, Latrunculin B, led to mitochondrial elongation in untreated Xenopus laevis melanocytes ^15^ and U2OS cells ^91^. While the direct link between Rac1, the actin cytoskeleton, and mitochondrial morphology/function remains to be mechanistically established in skeletal muscle, Rac1 preserves oxidative capacity by promoting actin cytoskeleton formation around damaged mitochondria in response to ischemia-induced acute injury in kidney proximal tubule epithelium ^94^. Additionally, targeting miR-142-3p, which down-regulates Rac1, impaired mitochondrial biogenesis and function in neuronal cells ^95^, while activated Rac1 increased mitochondria motility in MFT-16 ^96^. Together, these cell studies suggest that Rac1, via its role in actin cytoskeleton regulation, modulates mitochondrial morphology and respiration. This agrees with the enlarged subsarcolemmal and intermyofibrillar mitochondria with altered cristae density and partial mitochondrial damage observed in our study in mature skeletal muscle tissues. Moreover, the Rac1-deficiency-induced mitochondrial deterioration of oxidative function supports the growing evidence for mitochondrial dysfunction as a driver of sarcopenia ^3,97^. It is thus likely that Rac1-dependent mitochondrial dysfunction, in turn, resulted in premature muscle wasting in the Rac1 imKO mice, although causality was not established. In alignment, the deficiency of Rac1’s downstream targets, PAK1/2, contributed to age-related muscle wasting concomitant with the development of megaconial mitochondria ^13^. Altogether, these data indicate a new role of Rac1 in muscle mitochondrial biology, which, when lost, could accelerate muscle wasting with age.

Metabolomic and functional analyses implied impaired fatty acid oxidation and lipid handling in Rac1-deficient muscle, in line with the reduced mitochondrial respiration capacity. These findings establish a previously unrecognised link between Rac1 and fatty acid oxidation, uncovering a novel role for Rac1 in metabolic reprogramming of skeletal muscle. Studies in non-muscle systems implicate Rho GTPases in the regulation of the Hippo pathway, through modulation of YAP activity ^98–100^. In mature skeletal muscle, targeted disruption of YAP leads to incomplete fatty acid oxidation, lipotoxicity, and muscle wasting ^101,102^, raising the possibility that Rac1 may influence fatty acid metabolism via previously unrecognised Hippo-YAP signalling mechanisms. Moreover, in yeast *S. cerevisiae*, the Rac1 actin cytoskeleton-regulating downstream target, Cofilin, modified fatty acid metabolism via the mitochondrial outer membrane protein VDAC by altering actin dynamics ^19^. These findings align with our observation of blunted fatty acid oxidation, lowered palmitate uptake, and blocked TG usage during muscle contraction in Rac1 imKO mice, uncovering a previously unrecognised role for Rac1 in regulating metabolic flexibility and fatty acid utilisation. Similar observations have been made in other models of altered mitochondrial function. For example, inducible muscle-specific AMPKα1/α2 double KO mice also show reduced skeletal muscle contraction-stimulated palmitate oxidation ^103^, similar to the present results. Interestingly, this phenotype also mimics preclinical ^104^ and clinical ^105^ observations that muscle contraction-stimulated fatty acid oxidation capacity can be blunted in obesity. As a potential consequence of the blunted fatty acid oxidation, TG accumulated in Rac1 imKO muscle. In alignment, loss of Rac1 induced lipid accumulation in human hepatocyte Huh7 cells ^106^ and in response to ischemia-induced acute injury in the kidney proximal tubule epithelium ^94^. Interestingly, mitochondrial architecture has been suggested to influence fatty acid oxidation as forced mitochondrial elongation induced by Mfn2 overexpression, dominant negative DRP1 or siRNA-mediated DRP1 knockdown reduced long-chain fatty acid oxidation in primary hepatocytes ^107^, indicating a mechanism by which Rac1-dependent mitochondrial dysfunction could impair fatty acid oxidation and cause TG accumulation.

Finally, human sarcopenic muscle biopsies exhibited elevated Rac1 protein content, and Rac1 SNP variants were associated with muscle-wasting diseases and hyperlipidaemia, implicating Rac1 in the pathogenesis of sarcopenia and lipid metabolic diseases. In ageing human skeletal muscle, muscle energetics is associated with muscle quality ^6^, which in a longitudinal cohort of people above the age of 60 years predicted subsequent declines in muscle strength and lean mass ^108^. Additionally, the capacity for fatty acid oxidation during exercise is lower in elderly than in young subjects ^109–111^ as a consequence of diminished physical activity, decreased fat-free mass or ageing per se. Increased Rac1 in sarcopenic muscle likely compensates for declining mitochondrial efficiency, as Rac1 ablation in mice leads to premature muscle wasting. Our data indicate that Rac1 shifts from a homeostatic role in young muscle to a compensatory stress response in sarcopenia, where it acts to preserve energy metabolism and limit muscle atrophy.

Our findings support a model in which skeletal muscle Rac1 protein content increases with age to counteract sarcopenia through a hierarchical cascade. In this axis, Rac1-dependent actin remodelling preserves mitochondrial morphology and respiratory capacity, thereby maintaining fatty acid oxidative capacity and ultimately sustaining muscle mass (Fig. 8B). The precise mechanisms by which Rac1 protects against muscle decline – potentially through regulation of mitochondrial content and function, as well as fatty acid metabolism – and how these processes link Rac1 to muscle wasting were not mechanistically resolved in the present study. Mirroring observations in non-muscle cells, we found that skeletal muscle Rac1 localises to the mitochondria, where it potentially interacts with Bcl-2 to modulate mitophagy ^72^. Further evidence for a role in mitophagy comes from Rac1-deficient muscle, which showed PINK1 upregulation and increased pULK1 S757 levels. These findings, congruent with an accumulation of defective mitochondria, suggest a disruption in autophagy/mitophagy pathways. This is particular relevant as other proteostatic mechanisms, such as chaperone-mediated autophagy, are known to decline in aging mouse and human muscle ^112^. Moreover, genetically modified mice with deletion of chaperone-mediated autophagy in skeletal muscle exhibit reduced muscle strength and myofibre damage concomitant with mitochondrial dysfunction and defective SERCA function ^112^, potentially implicating calcium handling in the Rac1-dependent mitochondrial dysfunction, dysfunctional fatty acid metabolism and muscle wasting. Whether muscle Rac1 deficiency leads to defective SERCA function remains to be investigated. It also remains an intriguing unresolved question whether targeting Rac1 signalling (or its downstream effectors such as PAKs, actin regulators, or ROS pathways – all of which are suggested to be involved in mitochondrial regulation ^13,14,92,93,113^) could serve as a therapeutic avenue for muscle ageing.

In conclusion, using genetically modified mice, multi-omics analyses, and functional assays, we show that loss of Rac1 contributes to alterations in mitochondrial morphology and respiration, as well as fatty acid uptake and metabolism, contributing to muscle decline with age. Notably, human sarcopenic muscle biopsies exhibited elevated Rac1 protein content, which negatively correlated with multiple mitochondrial respiratory complexes, and Rac1 SNPs were linked to lipid metabolic disorders and muscle-wasting diseases, highlighting translational relevance. Together, these data position Rac1 as a previously unrecognised modulator of mitochondrial integrity and fatty acid metabolism, potentially with implications for sarcopenia and other muscle-wasting conditions. Targeting this pathway may offer therapeutic avenues to improve muscle function in ageing and mitochondrial dysfunction–associated muscle loss.

## Acknowledgements

We thank our colleagues at the August Krogh Section for Human and Molecular Physiology, Department of Nutrition, Exercise, and Sports (NEXS), Faculty of Science, University of Copenhagen (UCPH), Denmark (DK), and Division of Translational Metabolism, Department of Biomedical Sciences (BMI), Faculty of Health and Medical Sciences, UCPH, DK, for fruitful discussions on this topic. We acknowledge the skilled technical assistance of Irene B. Nielsen, Jacob F. Jeppesen (NEXS, Faculty of Science, UCPH, DK), Nanna J. Hahn, Mona S. Ali, Michala Carlsson, Katharina Stohlmann (BMI, Faculty of Health and Medical Sciences, UCPH, DK), and Peter Schjerling (Institute of Sports Medicine Copenhagen, Bispebjerg Hospital, DK). Mouse experiments were performed at the Animal Core Facility, Faculty of Health and Medical Sciences, UCPH, DK. We acknowledge the Histolab, Faculty of Health and Medical Sciences, UCPH, DK. Imaging by transmission electron microscopy was performed at the Core Facility for Integrated Bioimaging, Faculty of Health and Medical Sciences, UCPH, DK. Mass spectrometry-based proteomic analyses on muscles from middle-aged male Rac1 imKO mice were performed by the Proteomics Research Infrastructure (PRI) at UCPH, supported by the Novo Nordisk Foundation (NNF) (grant agreement number NNF19SA0059305). We want to acknowledge the participants and investigators of the FinnGen study. We thank Professor D. Grahame Hardie (University of Dundee, Scotland, UK) for the kind donation of the PDH-1Eα and PDK4 antibodies for this study. Illustrations were generated using https://Biorender.com (RRID:SCR_018361). Graphs were generated with GraphPad PRISM (RRID:SCR_002798) or R (RRID:SCR_001905) and the figures assembled in Inkscape (RRID:SCR_014479).

This study was supported by the Lundbeck Foundation (PhD fellowship, R208-2015-3388 and postdoctoral fellowship, R322-2019-2688 to LLVM), the Novo Nordisk Foundation (NNF20OC0063709 to EAR; NNF16OC0023418 and NNF18OC0032082 to LS), Independent Research Fund Denmark (4004-00233B and 9039-00170B to LS), the Carlsberg Foundation (CF21-0369 and CF24-1408 to LS), and the EFSD/Novo Nordisk Foundation Future Leaders Award (NNF24SA0094136 to LS).

## Author Contributions

Conceptualisation: LLVM, EAR, LS. Formal analysis: LLVM, ZKJO, JM, BLP. Funding Acquisition: LLVM, EAR, LS. Investigation: LLVM, SHR, EF, ABJ, NRA, AG, JLB, ENP, JM, SAN, BCB, NØ, HP, BK, DJ, BLP, JN, SL, EAR, LS. Methodology: LLVM, ABJ, TCPP, SAN, BCB, BK, BLP, JN SL, EAR, LS. Project administration: LLVM, LS. Resources: TCPP, AK, JA, MK, HP. Supervision: BCB, NØ, MK, HP, BK, DJ, SL, EAR, LS. Visualisation: LLVM. Writing – Original Draft Preparation: LLVM, LS. Writing – Review & Editing: LLVM, SHR, EF, ABJ, NRA, AG, ZKJO, TCPP, JLB, ENP, JM, AK, JA, SAN, BCB, NØ, MK, HP, BK, DJ, BLP, JN, SL, EAR, LS.

## Data and Resource Availability

The authors confirm that the data supporting the findings of this study are available within the article and/or supporting information. The mass spectrometry-based proteomic dataset from adult Rac1 imKO mice is being deposited to the ProteomeXchange Consortium, and the dataset identifier will be provided to the reviewers. The mass spectrometry proteomics data from middle-aged Rac1 imKO mice have been deposited to the ProteomeXchange Consortium (http://proteomecentral.proteomexchange.org) via the PRIDE partner repository with the data set identifier PXD079634 (https://www.ebi.ac.uk/pride/archive/projects/). The following publicly available GEO dataset was analysed in the present study: GSE145480.

---

**Figure S1:**
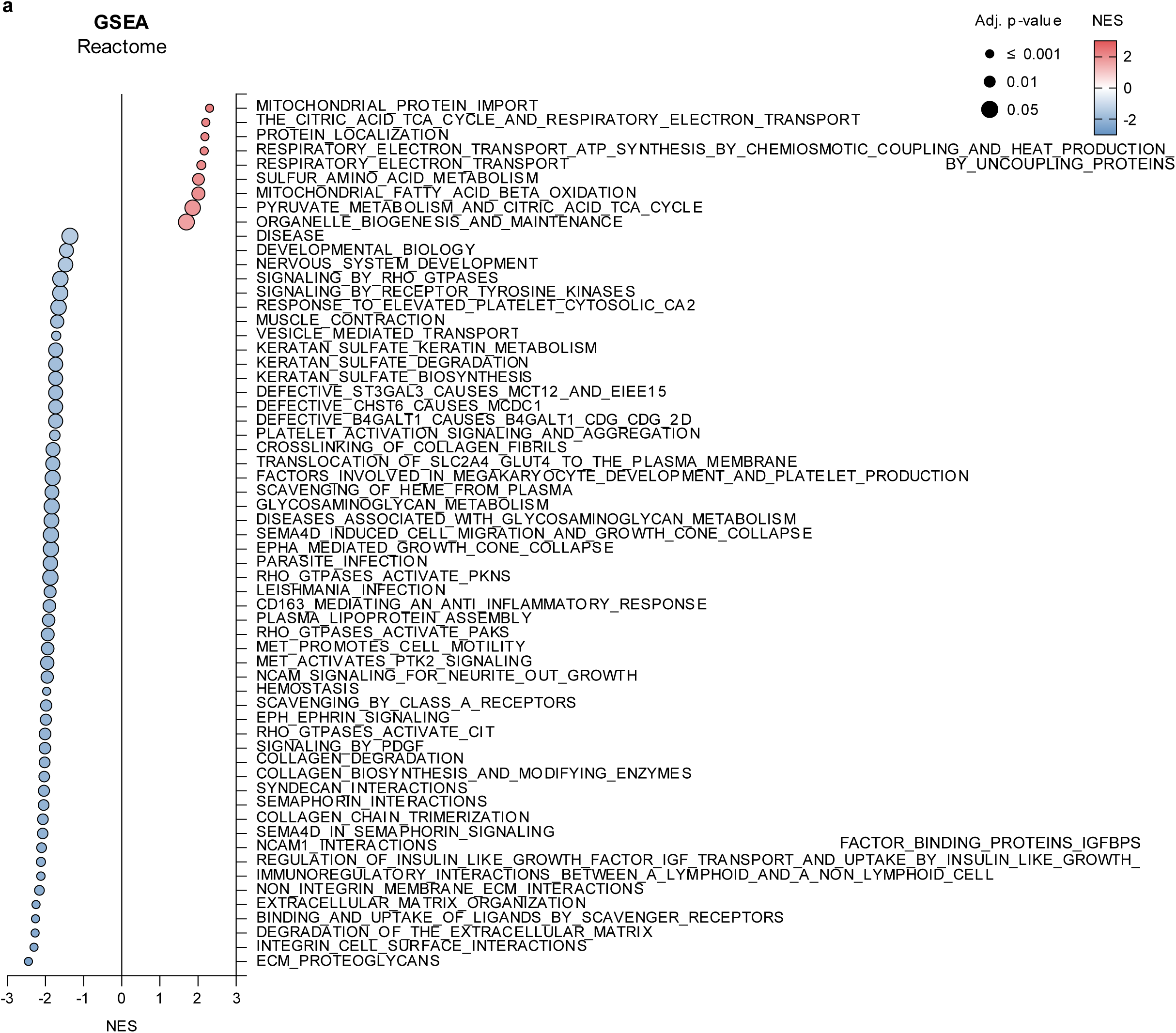
(A) Reactome Gene Set Enrichment Analysis (GSEA) showing Normalised Enrichment Score (NES) values for each statistically significant gene set and adjusted p-values in gastrocnemius muscle from adult inducible muscle-specific Rac1 knockout (imKO) mice or littermate controls. *n = 4/1* (male/female).

**Figure S2:**
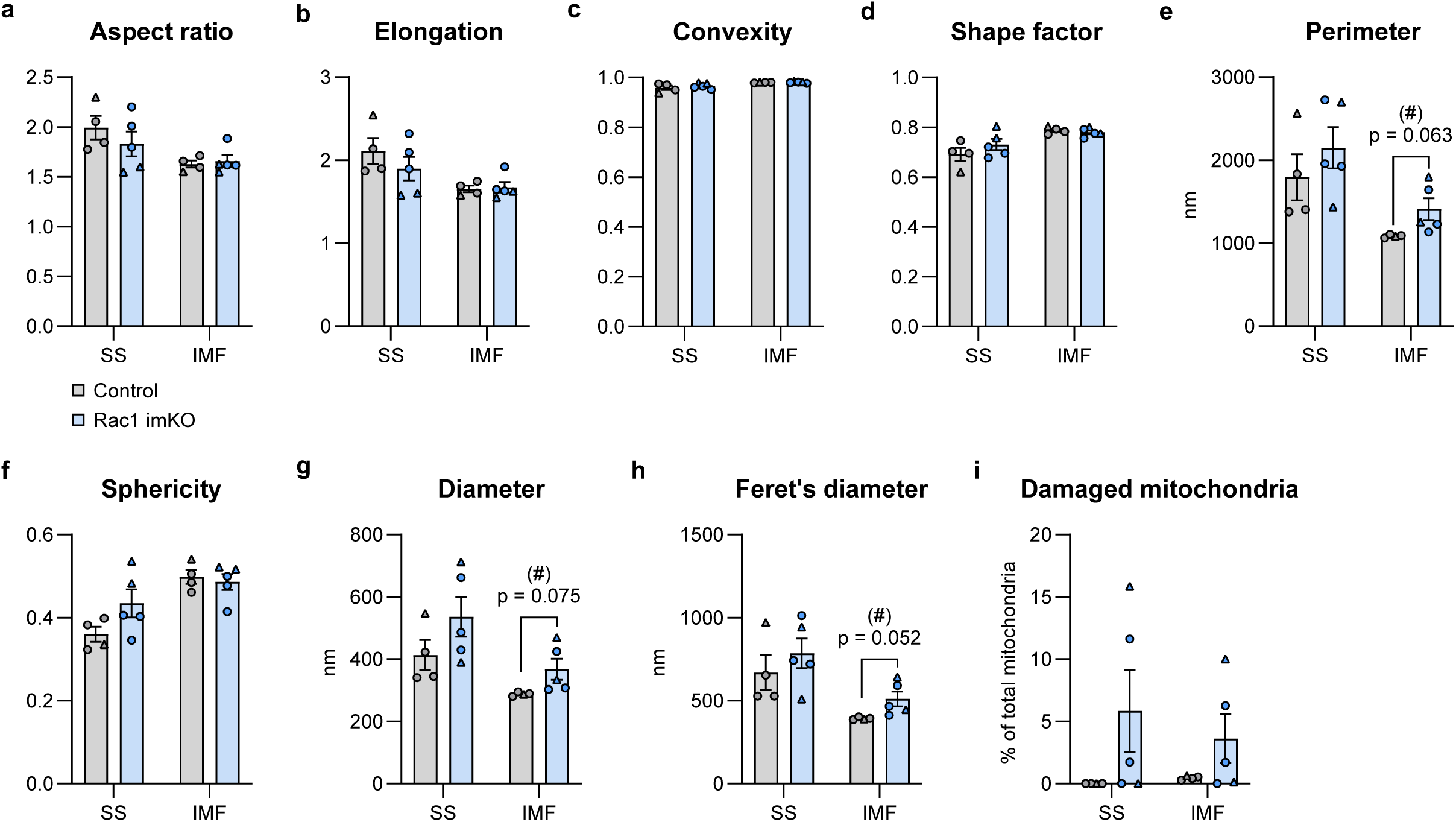
Subsarcolemmal (SS) and intermyofibrillar (IMF) mitochondrial (A) aspect ratio, (B) elongation, (C) convexity, (D) shape factor, (E) perimeter, (F) sphericity, (G) mean inner diameter, (H) maximal ferret’s diameter, and (I) percentage of damaged mitochondria in quadriceps muscle from inducible muscle-specific Rac1 knockout (imKO) mice or littermate controls. Control, *n = 3/1* (male/female); Rac1 imKO, *n = 3/2*. Data were evaluated with a Student’s t-test: Effect of Rac1 imKO (#) (p < 0.1). Data are presented as mean ± SEM with individual data points shown. ◯ Male, △ Female.

**Figure S3:**
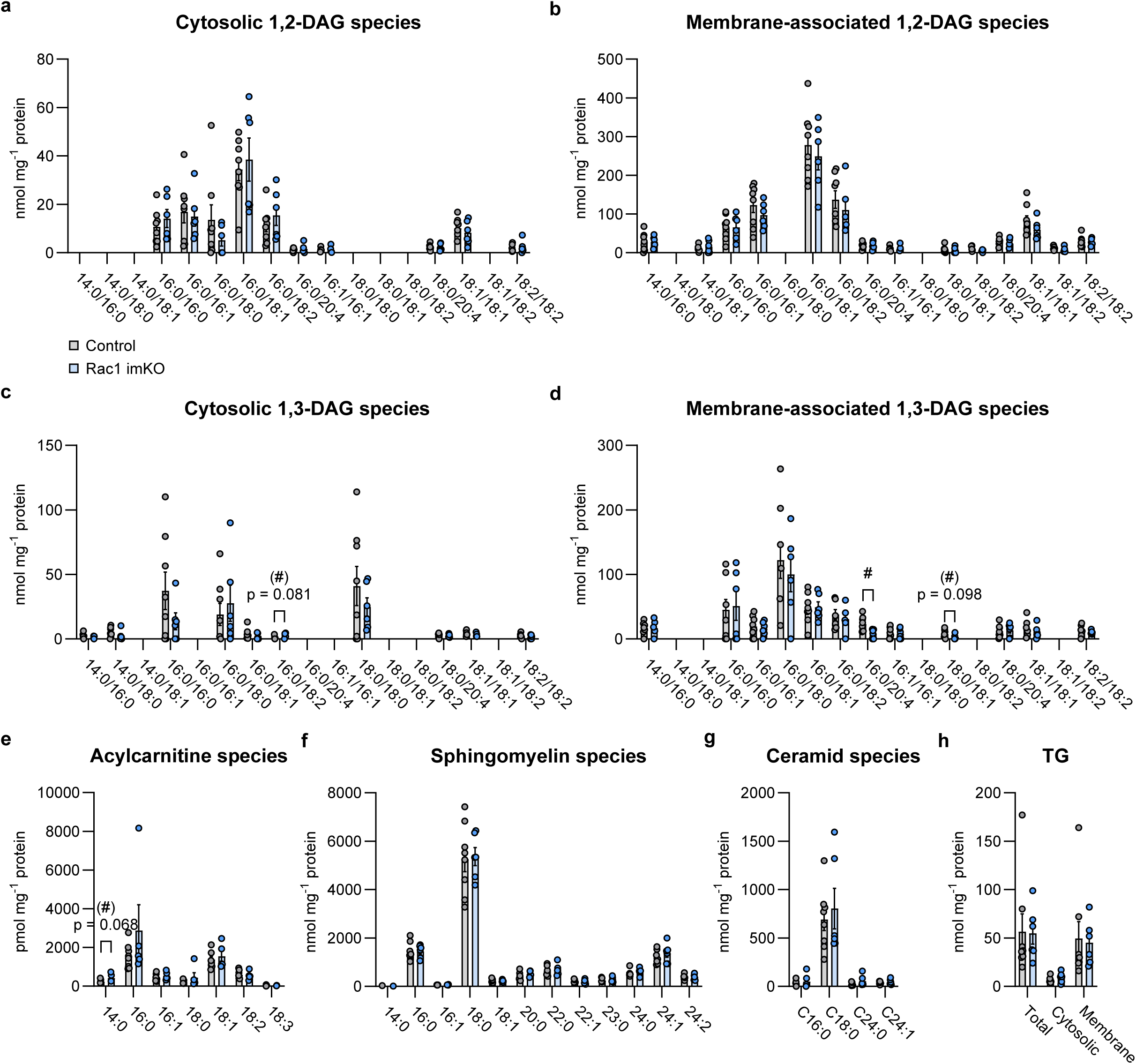
(A) Cytosolic 1,2-diacylglycerol (DAG), (B) membrane-associated 1,2-DAG, (C) cytosolic 1,3-DAG, (D) membrane-associated 1,3-DAG, (E) acylcarnitine, (F) sphingomyelin, (G) ceramide, and (H) triacylglycerol (TG) species in quadriceps muscle from male inducible muscle-specific Rac1 knockout (imKO) mice or littermate controls. *n = 8/6* (Control/Rac1 imKO). Data on acylcarnitine species were not obtained for *n = 1* Rac1 imKO mouse. Data were evaluated with a Student’s t-test. Significant Student’s t-test: Effect of Rac1 imKO (#)/# (p < 0.1/0.05). Data are presented as mean ± SEM with individual data points shown.

**Figure S4:**
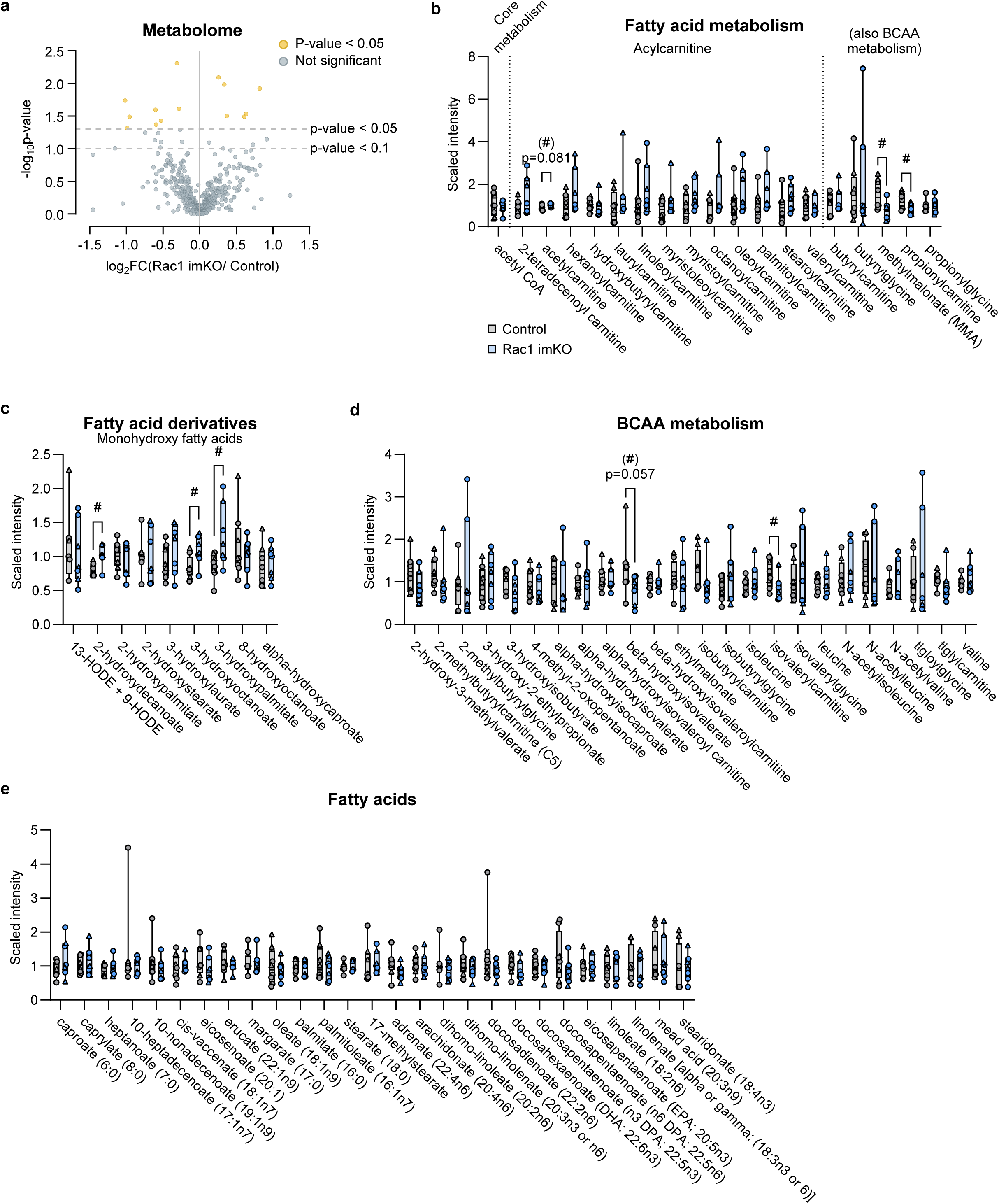
(A) Volcano plot of all metabolites detected with Ultrahigh Performance Liquid Chromatography-Tandem Mass Spectroscopy (UPLC-MS/MS) in gastrocnemius muscle from adult inducible muscle-specific Rac1 knockout (imKO) mice or littermate controls in response to acute treadmill running at 70-75% of the maximal running speed for 30 minutes (10° incline). Control, *n = 5/3* (male/female); Rac1 imKO, *n = 4/3*. (B) Fatty acid metabolism, (C) fatty acid derivatives, (D) branched chain amino acid (BCAA) metabolism and (E) fatty acid metabolites. Data were evaluated with a Welch’s t-test: Effect of Rac1 imKO (#)/# (p < 0.1/0.05). Data are presented as mean ± SEM with individual data points shown. ◯ Male, △ Female.

**Figure S5:**
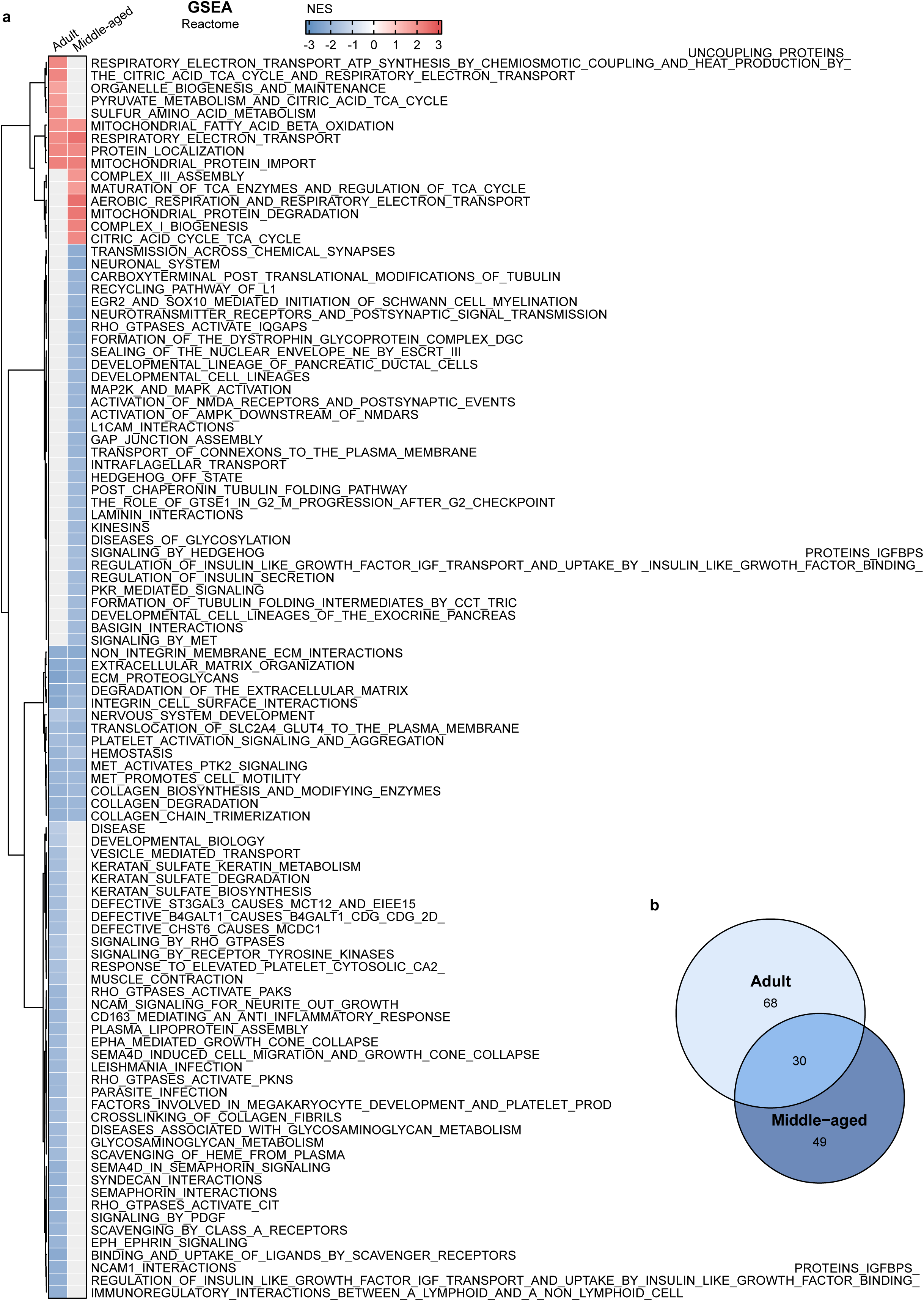
(A) Reactome Gene Set Enrichment Analysis (GSEA) showing Normalised Enrichment Score (NES) values for each statistically significant gene set in gastrocnemius muscle from middle-aged male inducible muscle-specific Rac1 knockout (imKO) mice or littermate controls. (B) Overlapping gene sets significantly enriched or depleted in GSEA with changes in the proteome of adult and middle-aged Rac1 imKO mice compared to control. GSEA was conducted separately for the proteome in adult and middle-aged mice.

**Figure S6:**
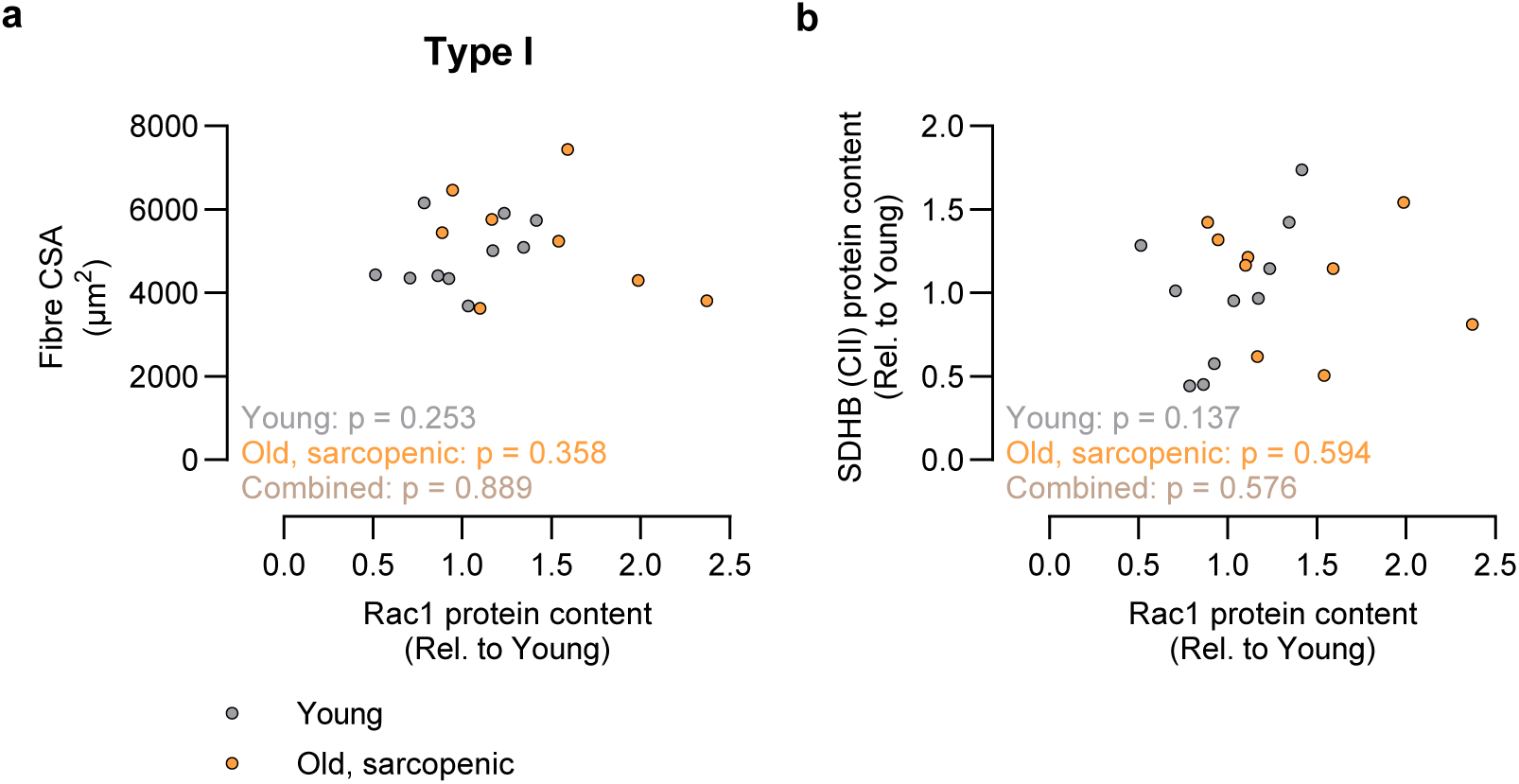
(A) Correlation between Rac1 protein content and type I fibre cross-sectional area (CSA; type I fibre CSA was previously published ^28,30^) and (B) OXPHOS CII subunit SDHB protein content in old, sarcopenic vastus lateralis muscle compared to young control. *n = 10/9* (Young/Old, sarcopenic). For type I CSA, data were not obtained for *n = 1*. Data were evaluated with Pearson’s correlation.

**Table S1.**
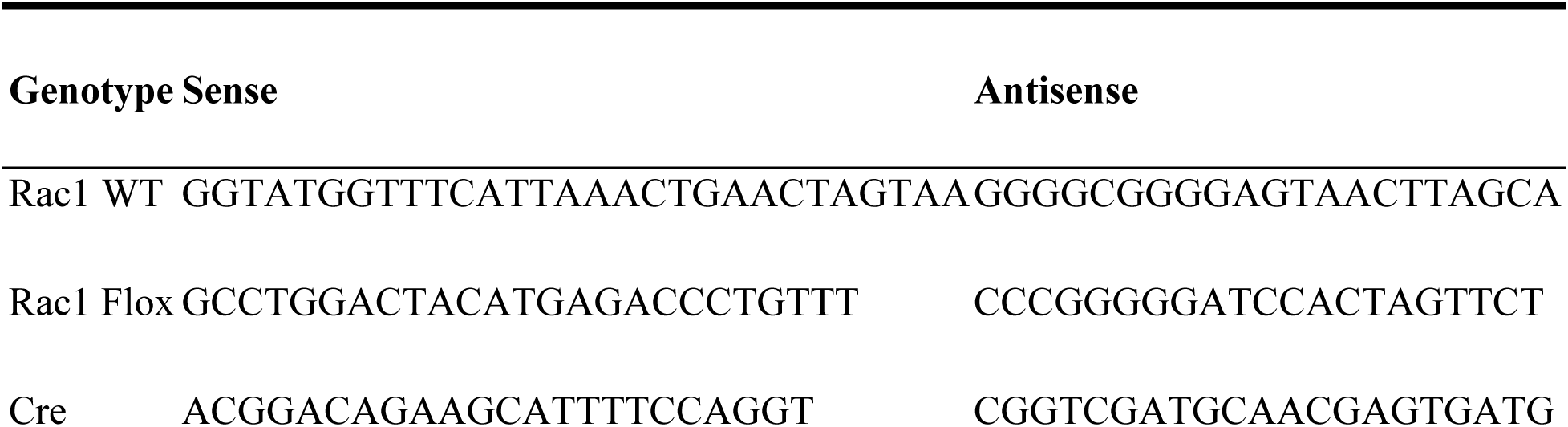
Primers for mouse genotyping.

**Table S2.**
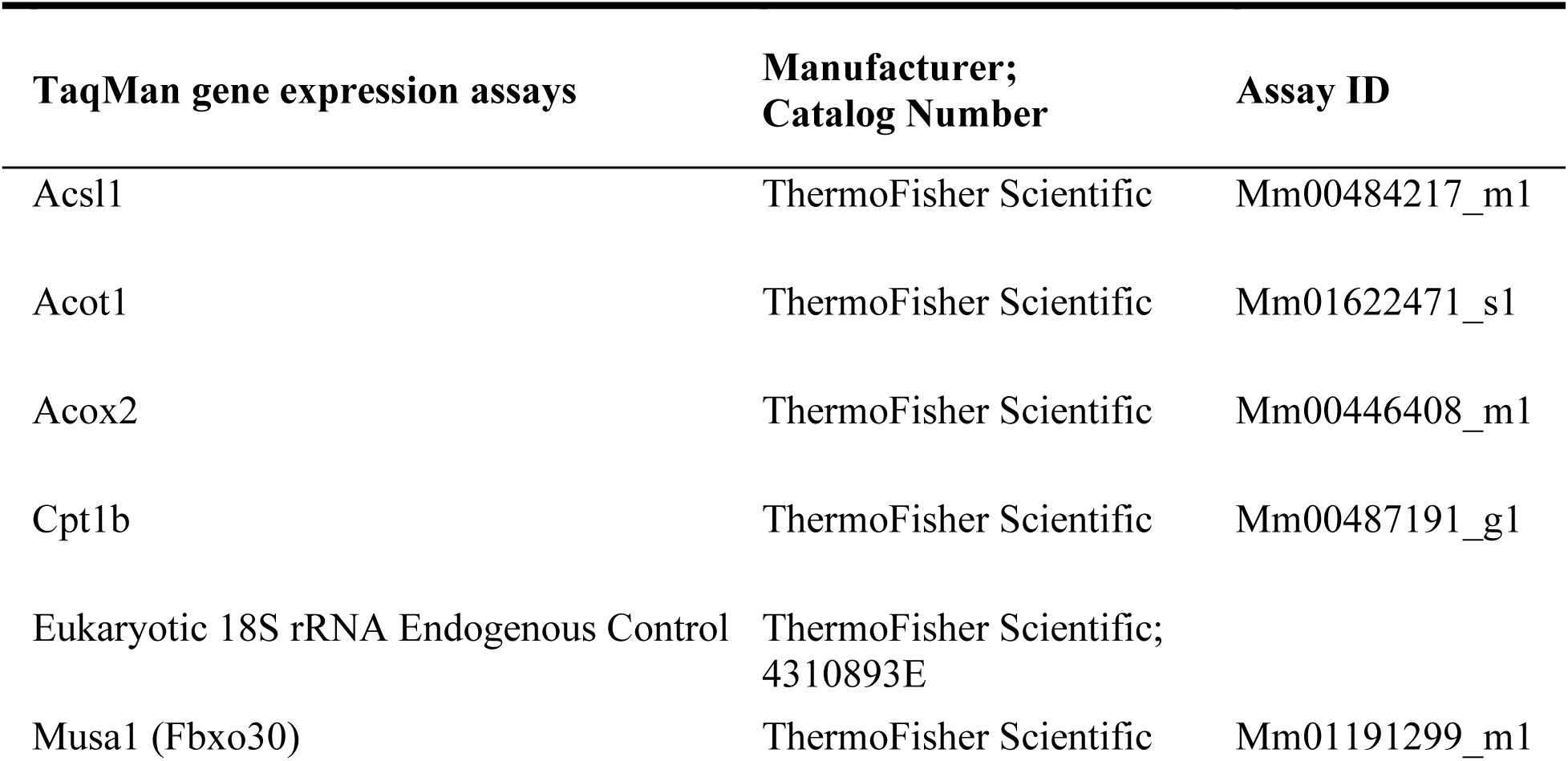
TaqMan gene expression assays.

**Table S3.**
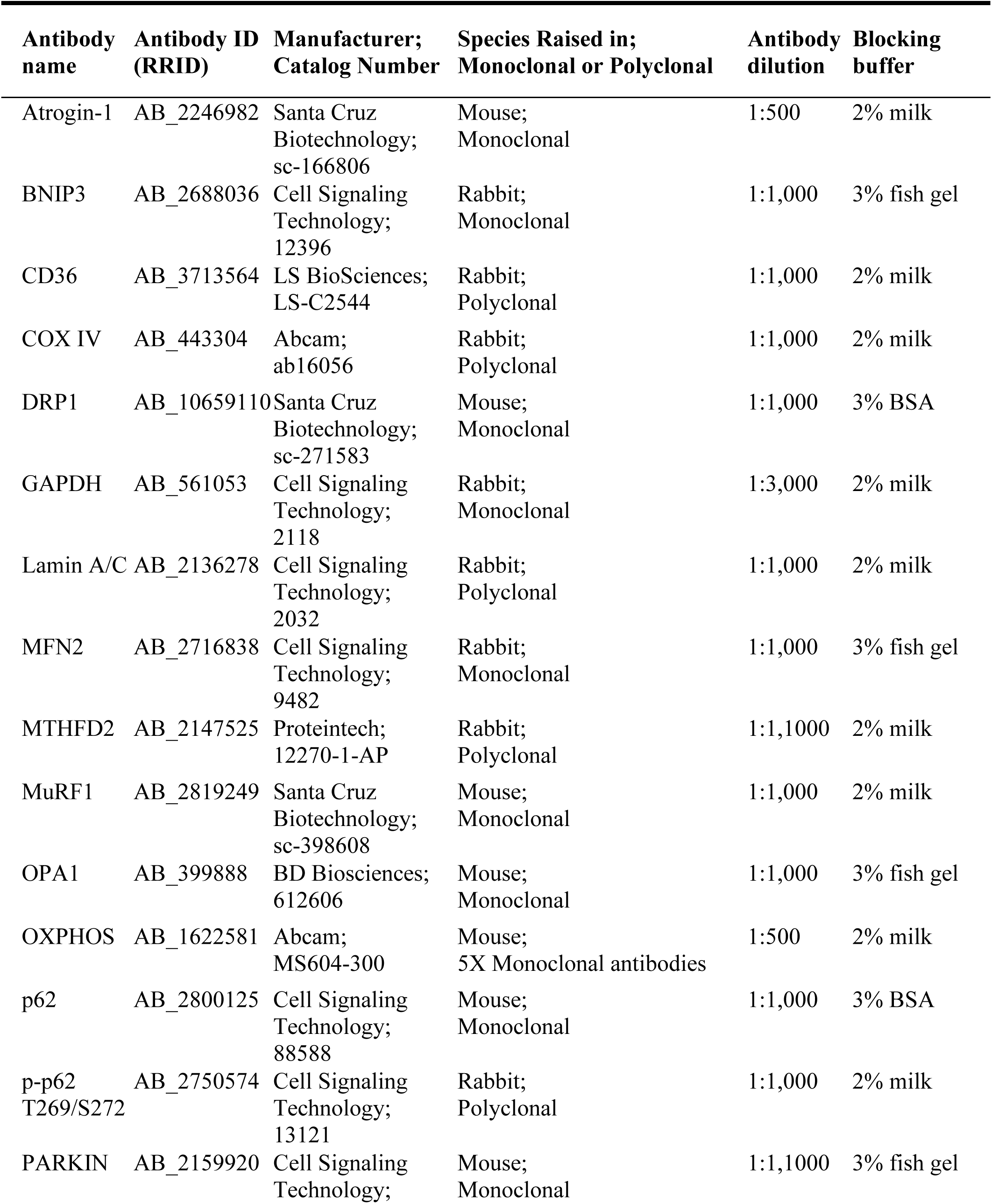
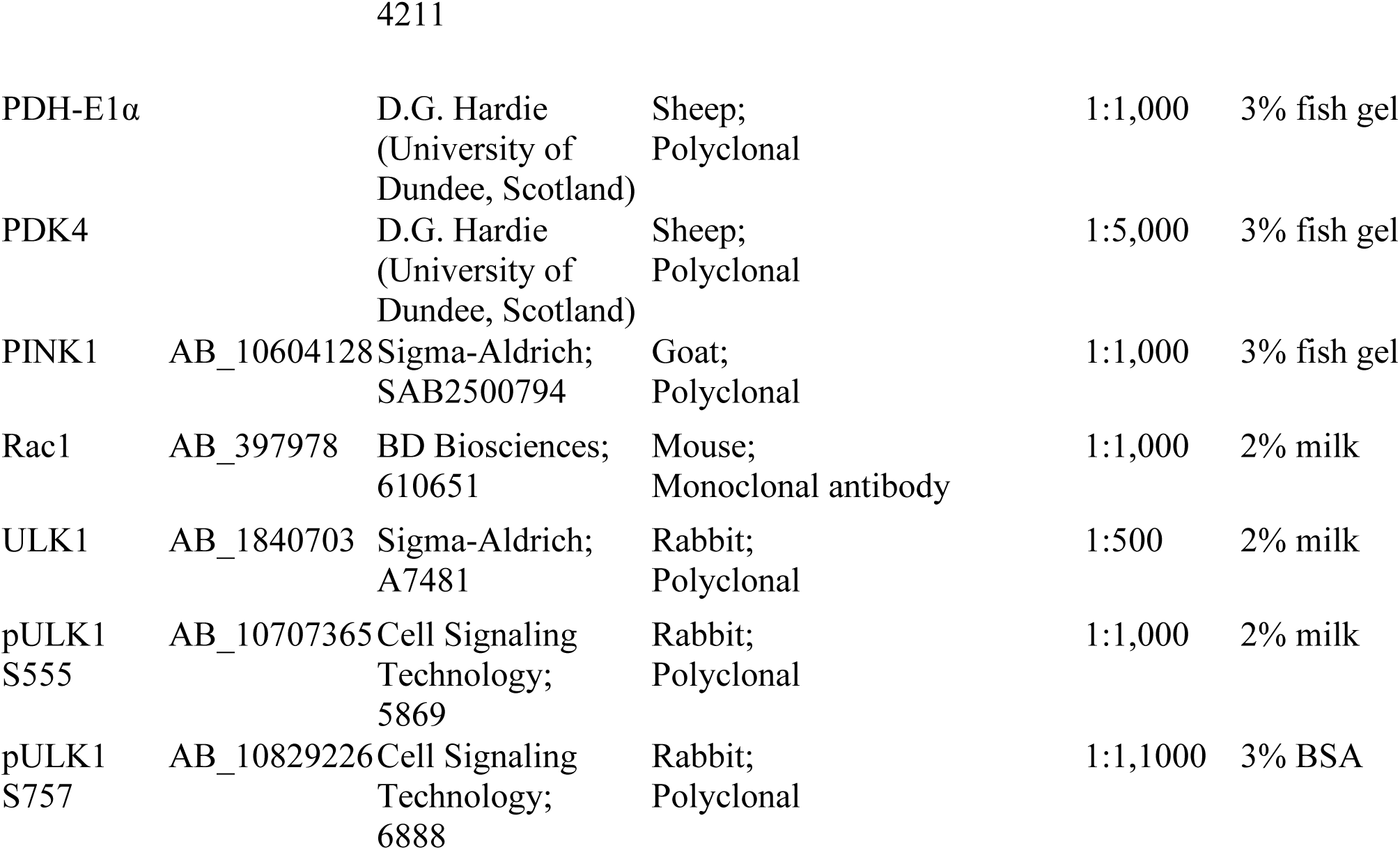
Antibodies.

